# Active intermixing of indirect and direct neurons builds the striatal mosaic

**DOI:** 10.1101/378372

**Authors:** Andrea Tinterri, Fabien Menardy, Marco A. Diana, Ludmilla Lokmane, Maryama Keita, Fanny Coulpier, Sophie Lemoine, Caroline Mailhes, Benjamin Mathieu, Paloma Merchan-Sala, Kenneth Campbell, Ildiko Gyory, Rudolf Grosschedl, Daniela Popa, Sonia Garel

## Abstract

The striatum controls behaviors via the activity of direct and indirect pathway projection neurons (dSPN and iSPN) that are intermingled in all compartments. While such mosaic ensures the balanced activity of the two pathways, how it emerges remains largely unknown. Here, we show that both SPN populations are specified embryonically and progressively intermix through multidirectional iSPN migration. Using conditional mutants of the dSPN-specific transcription factor Ebf1, we found that inactivating this gene impaired selective dSPN properties, including axon pathfinding, whereas molecular and functional features of iSPN were preserved. Remarkably, *Ebf1* mutation disrupted iSPN/dSPN intermixing, resulting in an uneven distribution. Such architectural defect was selective of the matrix compartment, revealing that intermixing is a parallel process to compartment formation. Our study reveals that, while iSPN/dSPN specification is largely independent, their intermingling emerges from an active migration of iSPN, thereby providing a novel framework for the building of striatal architecture.

## Introduction

The dorsal striatum controls major brain functions, such as motor behaviors and habit formation, through the coordinated activation of descending direct and indirect pathways. Consistently, loss or damage of striatal projection neurons (SPN) is associated with a spectrum of pathologies that include Parkinson’s disease, Huntington’s disease and obsessive-compulsive disorders. SPN are medium-spiny GABAergic neurons which account for 90-95% of striatal neurons and are divided into two subtypes: i) direct SPN (dSPN) project to the substantia nigra (SN) and express the D1 dopamine receptor and the neuropeptide Substance P; (ii) indirect SPN (iSPN) send axons to the globus pallidus (GP) and express D2 dopamine receptor and the opioid peptide Enkephalin. Functionally, it is proposed that dSPN activation promotes action selection and positive reinforcement, whereas iSPN are important to suppress unwanted motor sequences^1–7^. dSPN and iSPN are completely intermixed within the striatum, thereby forming a relatively uniform mosaic^8–10^. Building on this relatively simple cellular organization, the dorsal striatum is split into at least two major compartments, cellular islands known as striosomes and a surrounding matrix, which form immunohistologically and functionally distinct modules differing by their input and output^11–17^. In such precise three-dimensional organization, the intermingling of dSPN and iSPN ensures the balanced activation of direct and indirect pathways.

While functional studies revealed distinct roles for dSPN/iSPN and striosomes/matrix, much less is known on how they develop. SPN are generated in the embryonic lateral ganglionic eminence (LGE) and migrate radially to form the striatum^18–24^. Both SPN subtypes are present in the entire striatum, except for a restricted caudal domain^8^, and several transcription factors have been involved in their generation and differentiation including Ctip2, FoxP1 and FoxP2 ^25–40^. In addition, distinct transcriptomic programs have been involved in the specification of either dSPN or iSPN, including Islet1 and Ebf1 for dSPN or Sp9 and Six3 for iSPN^23, 41–46^. In particular, the conditional inactivation of *Islet1* impairs the differentiation of early-born dSPN^41, 45^ and the full inactivation of *Ebf1* leads to a defective development of late-born dSPN, disorganized projections to the SN and altered survival of matrix dSPN at postnatal stages^42–44^. Regardless of their subtype, early progenitors give rise to striosome SPNs whereas later-derived progenitors generate matrix SPN, which form the two compartments potentially via migration and selective cell-sorting^17, 24, 47–51^. In contrast, how dSPN and iSPN are specified and integrated into striatal circuits remains largely to be characterized.

Here we show that formation of the striatal mosaic relies on the early specification of SPN subtypes combined with a dSPN-dependent tangential migration of iSPN. We found that dSPN and iSPN form two molecularly-defined populations throughout development, which only progressively intersperse. Using time-lapse imaging, we observed that iSPN undergo multidirectional migration, thereby intermixing with dSPN. Furthermore, we took advantage of a unique combination of genetic tools, including embryonic *in vivo* fate map and conditional knockout models, to examine the role of *Ebf1* in the differentiation of dSPN and striatal mosaic formation. We found that *Ebf1* deletion in dSPN (i) perturbed specific aspects of direct neurons differentiation, leading to defective axogenesis and integration in cortico-striatal circuits, without affecting iSPN cardinal properties (ii) impaired intermixing of dSPN and iSPN in the matrix. These results establish *Ebf1* as a master regulator of dSPN connectivity and intermixing with iSPN in the matrix compartment. Furthermore, our study reveals that the active intermingling of early-specified dSPN and iSPN is required to assemble the striatal mosaic.

## Results

### Defining a molecular fingerprint of embryonic dorsal dSPN

To investigate how dSPN and iSPN are specified, positioned and assembled in the striatum, we searched for specific markers of these two neuronal populations during embryogenesis. To this aim, we took advantage of the BAC transgenic mouse line *Drd2-EGFP^52^*, which has been widely used to label iSPN in neonates and is expressed embryonically^53, 54^. In addition, we crossed *R26^mt/mt^* reporter mice^55^ with *Islet1^Cre^* transgenic mice, which mediate Cre-recombination in dSPN from embryonic stages^41, 45^. We first confirmed that these two distinct transgenic lines respectively labeled iSPN and dSPN at postnatal day (P) 5, when the two populations can be unambiguously defined^53, 56^. In *Islet1^Cre^;R26^mt/+^;Drd2-EGFP* mice, we found that tdTomato-positive (tom+) cells constituted approximately half of all Ctip2+ SPN^25^ in both striosome and matrix compartments (as defined in Fig. S1) and were largely unlabeled by *Drd2-EGFP* (Figs. 1a-d). Using these two mouse lines, we found that iSPN and dSPN lineages are non-overlapping throughout embryogenesis, consistently with previous studies focusing on SPN subtypes specification ^23, 41, 43–46^. We furthermore identified high expression of transcription factors FoxP2 and Ebf1 as specific of dSPN at E13.5 and E17.5 (Figs. 1e-h). These results identify a dSPN developmental fingerprint which, combined to the iSPN-specific *Drd2-EGFP* expression, delineates two segregated populations of Ctip2+/Foxp1+ SPN throughout development (Fig. 1i). Our findings thus indicate that dSPN and iSPN exhibit distinct molecular identities from the earliest steps of striatogenesis and provides us with tools to follow their integration in the striatal architecture.

**Figure 1.**
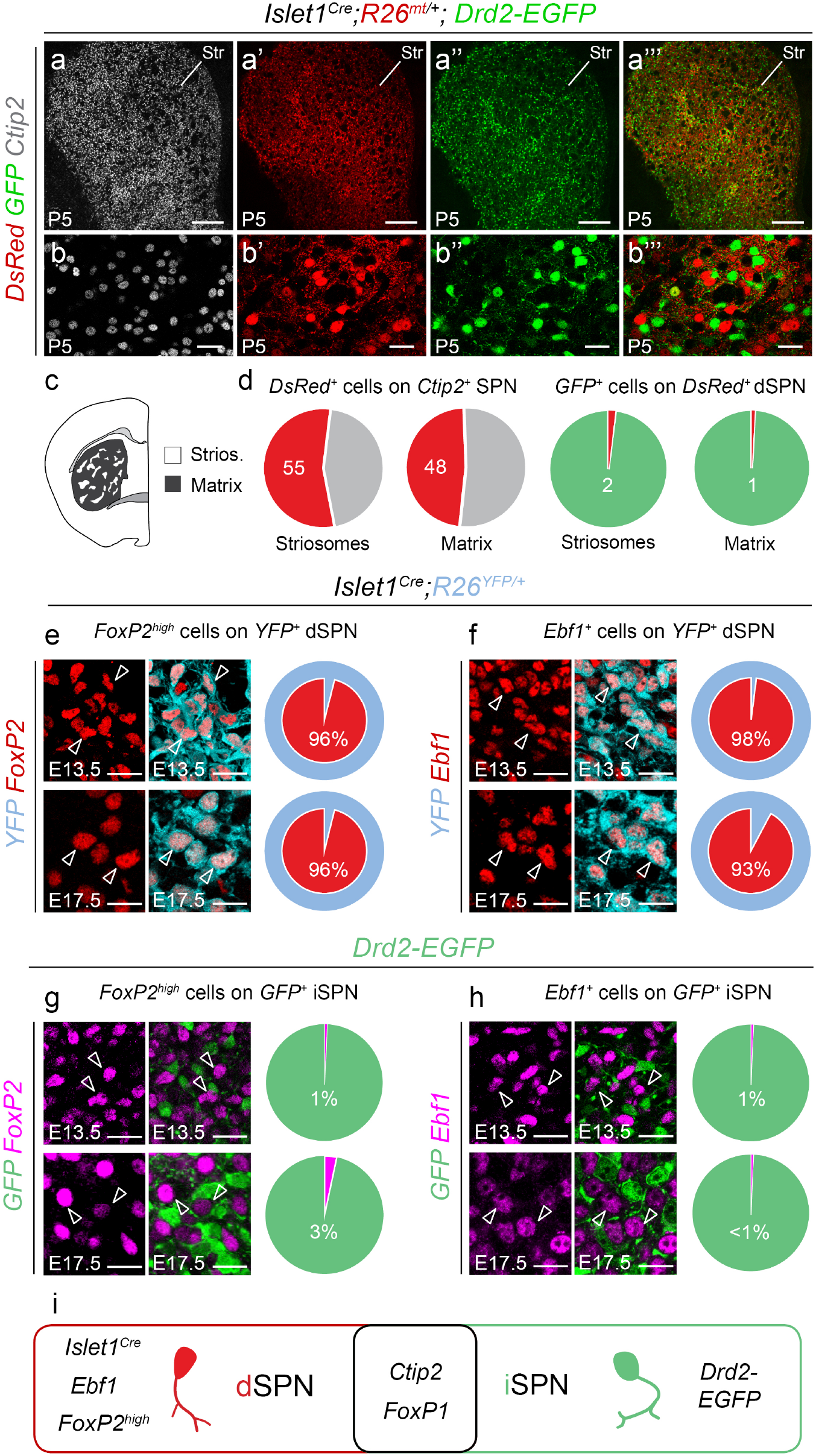
Definition of an embryonic molecular fingerprint of dSPN. (**a-a**”’) Coronal section of P5 *Islet1^Cre^;R26^mt/+^;Drd2-EGFP* mice showing Ctip2-immunostaining in all SPN (a) and non-overlapping labelling of tdTomato in dSPN and GFP in iSPN (a’-a”) in the dorsal striatum (Str). (**b-b’”**) Close-up highlighting the overall lack of overlap between tdTomato and GFP labelings. **(c)** Schematic representation of striatal compartments. **(d)** Quantification of the percentage of tdTomato+ dSPN in striatal compartments (n=3 mice), on the total of Ctip2+ cells (55±2% in Strios., 48±7% in matrix) and on the total of GFP+ cells (2±1% in Strios., 1±1% in matrix). (**e-f**) Co-immunostaining for FoxP2^high^ and Ebf1 labels the YFP-positive dSPN lineage in *Islet1^Cre^;R26^YFP/+^* embryos at both E13.5 (96±1% for Foxp2^high^; 98±1%for Ebf1) and E17.5 (96±2% for Foxp2^high^; 93%±1% for Ebf1) (n=3 mice for each stage). (**g-h**) Conversely, *FoxP2* and *Ebf1* are scarcely co-expressed by *Drd2-EGFP+* cells at E13.5 (1±1% and 1 ±0,1 % of GFP+ cells, respectively) and E17.5 (3±3% and 1 ±0,1 % of GFP+ cells, respectively)(n=3 mice for each stage). (**i**) Schematic representation of the embryonic genes common and specific to dSPN and iSPN. Results are presented as mean values ± sem. Scale bars equal 250 μm (a-a”’), and 25 μm (b-b”’, e-h). Str, striatum; Strios., Striosomes.

### iSPN progressively intermix with dSPN via intrastriatal migration

We took advantage of the early molecular fingerprints to monitor the dynamics of SPN distribution and thereby extend our comprehension of how the two populations are specified and assembled (Fig. 2). Previous studies showed that all SPN are generated in the LGE over an extended period of time and migrate radially to form the striatum^18–22, 24^. Moreover, birthdating experiments indicate that a majority of earliest-born SPN are dSPN^24, 48^. Consistently, we found that the striatum at E12.5 contained mostly dSPN neurons and few scattered GFP+ iSPN (Figs. 2a-a”’). GFP+ iSPN were detected in larger numbers at E13.5 and E14.5, albeit concentrated in the lateral striatum (Figs. 2b-c”), which contains early-born neurons that will mostly contribute to striosomes^20, 47^. From E15.5 onwards, we detected a progressively increasing number of GFP+ iSPN in the initially dSPN-dense part of the medial striatum (Figs. 2d-e”). Thus iSPN generated in the LGE proliferative zones intersperse into dSPN-dense territories, first in the lateral striatum and then medially, in a gradual process that spans several days (Fig. 2f). This phenomenon was temporally and spatially overlapping with the formation of striosome and matrix compartments^20, 21, 24, 49^.

**Figure 2.**
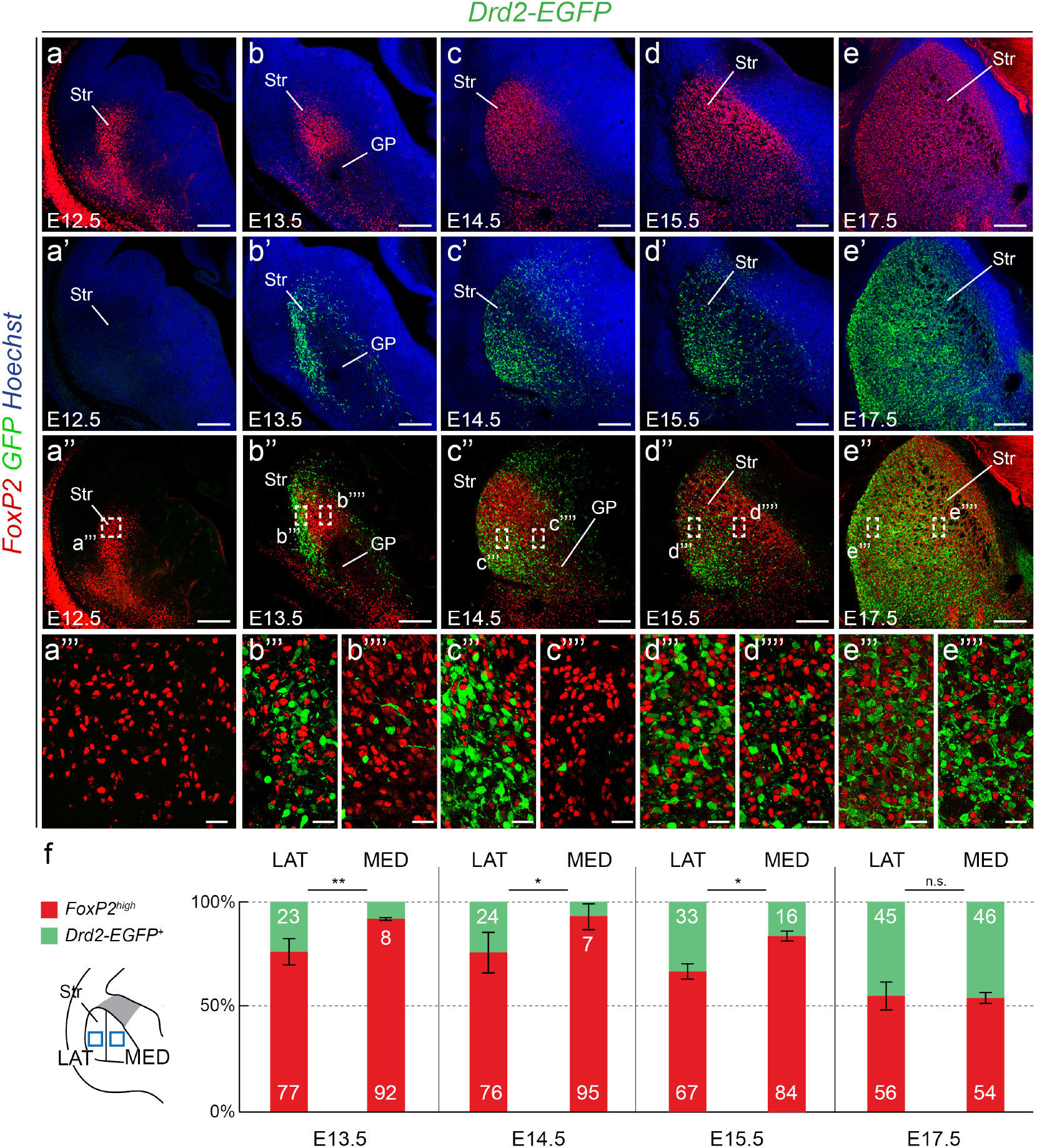
Progressive insertion of iSPN into the dSPN-dense striatal mantle. (**a-a”**) At E12.5, only FoxP2^+^ dSPN are located in the striatal mantle, while *Drd2-EGFP+* iSPN cannot be detected (inset in a”’). (**b-b”**) At E13.5, *Drd2-EGFP+* iSPN are mostly located in the lateral mantle (inset in b”’, 23±6% of total number of *Drd2-EGFP+* and *FoxP2+* cells) while almost none are detected medially (inset in b””, 8±1%, p=0.008). (**c-c”**) At E14.5, *Drd2-EGFP+* cells account for 24±10% of all SPN laterally, but only 7±6% medially (insets in c”’ and c””, p=0.02). (**d-d”**) At E15.5, more iSPN are found in the medial part of the striatum, but the distribution is still significantly different (insets in d”’ and d””, 33±3% laterally versus 16±4% medially, p=0.01). (**e-e”**) At E17.5, iSPN are distributed similarly in the lateral and medial striatum (insets in e”’ and e””, 45±7% laterally versus 46±2% medially, p>0.05), showing that iSPN progressively achieve a homogeneous distribution in the striatum. (f) Quantifications of the dSPN Foxp2^high^+/iSPN GFP+ distribution (at least n=3 mice for each stage). Results are presented as mean ± standard deviation. Scale bars equal 200 μm (a-e”) and 25 μm (a”‘-e””). GP, globus pallidus; LAT, lateral striatum; MED, medial striatum; Str, striatum.

Such progressive mosaic formation could be explained by: i) delayed expression of *Drd2-EGFP* compared to dSPN markers; ii) tangential dispersion of iSPN in the striatal mantle. To discriminate between these possibilities, we first examined whether immature iSPN in the LGE subventricular zone (SVZ) expressed *Drd2-EGFP* and whether timed EdU injections showed preferential labelling of either iSPN or dSPN. We found that immature iSPN already expressed *Drd2-EGFP* in the SVZ (Figs. S2a-b’) and that these neurons were produced continuously from E11 onwards (Figs. S2c-e), consistently with previous studies^24, 54^. iSPN however constitute a minority of the early-born SPN, as revealed by EdU staining (Figs. S2c-e) as well as labeling with the dSPN-specific Foxp2 immunostaining and *Islet1^Cre^* driven recombination (Figs. S2f-h). Taken together, our results indicate that *Drd2-EGFP* labels iSPN as they differentiate and enter the striatum primordium.

It is thus unlikely that mosaic formation is solely due to either a delay in *Drd2-EGFP* expression or a sequential production of SPN subtypes. To examine whether iSPN disperse tangentially inside the striatum, we performed two-photon time-lapse imaging on *Drd2-EGFP* embryonic slices^57^. We found that GFP+ iSPN show multidirectional saltatory migration within the striatal mantle at E15.5 (Figs. 3a-e; Movie S1). By performing cell behavior analyses over several time-lapse acquisitions, we found that iSPN have a global displacement away from the SVZ but present multidirectional trajectories inside the striatum (Figs. 3f-i and S3). In particular, iSPN changed their migration direction over time (Movie S1). Importantly, we confirmed that *Drd2-GFP+* cells in slices are Ctip2+ SPN and that the processes of these neurons also present multiple orientations in fixed embryonic tissue (Fig. S3). Thus, in contrast to the assumption that SPN only migrate radially, iSPN undergo a tangential migration within the striatum and intermix with dSPN.

**Figure 3.**
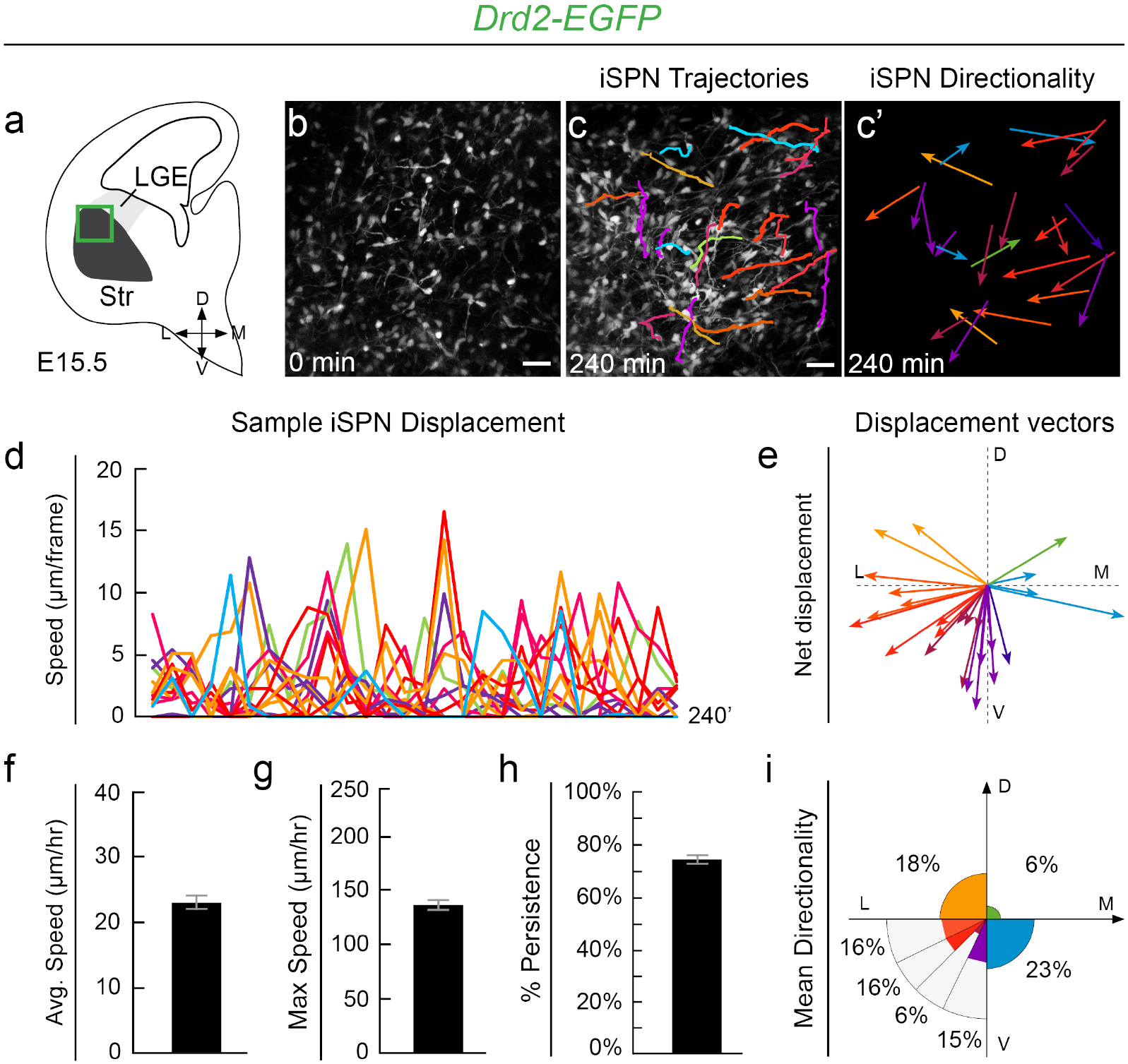
Tangential migration of iSPN in the striatal mantle. (**a**) Schematic coronal hemisection of a *Drd2-EGFP+* slice illustrating the area where multi-photon time-lapse imaging was performed. (**b-e**) Sample image (b, c with superimposed trajectories of individual cells) from a video tracking of iSPN migration allowing the identification of cell directionality (**c’**), saltatory behavior (d) and multidirectional displacement vectors (e; colors illustrate directionality). (**f-i**) Analysis of all tracked iSPN (n=61 cells, n=4 independent experiments) shows a moderate average net speed (**f**, 21.3±1 μm/hr), a high maximum speed (g, 124.8±5 μm/hr) and strong persistence (h, 74±15%, ratio between net displacement speed and total distance traveled). (**i**) Analysis of mean directionality shows that iSPN migrate tangentially within the striatal mantle (color code similar to c and e). Results are presented as mean ± s.e.m. Scale bar equals 30 μm. D, dorsal; L, lateral; LGE, lateral ganglionic eminence; M, medial; Str, striatum; V, ventral.

Taken together, our work shows that dSPN and iSPN form segregated embryonic populations that intermix over time, first in the lateral early-born proto-striosomes and then in the medial region. This process occurs at least in part via a multidirectional migration of iSPN thereby raising questions about the underlying mechanisms and possible interactions between the two populations.

### *Ebf1* conditional deletion affects specific aspects of dSPN differentiation

To address these issues, we searched for mouse models selectively perturbing the development of one of the SPN subtypes. Transcription factors specifically involved in either dSPN or iSPN differentiation include Islet1 and Ebf1 or Sp9 and Six3, respectively^23, 41–46^. In contrast to *Islet1, Six3* or *Sp9* deletions, which induce cell death in the targeted population^23, 41, 45, 46^, *Ebf1* inactivation has been previously proposed to impair dSPN wiring in the striatal matrix^42–44^. We thus generated and compared two different conditional *Ebf1* mutants (cKO), using the dSPN-restricted *Islet1^Cre^* line and the *Dlx5/6::Cre.* Indeed, *Islet1^Cre^* drives recombination in dSPN and cholinergic interneurons^47, 51, 41, 45, 58–60^ and *Dlx5/6::Cre* recombines in all SPN and all interneurons^61–65^. Importantly, Ebf1 is not expressed in striatal interneurons (Figs. S1a-c) and the two cre lines are nonoverlapping outside of the ventral telencephalon^58, 60, 61, 65–68^. Thus comparing the phenotypes shared in these two conditional mutants (cKO) allowed us to determine the deficits due to *Ebf1* inactivation in dSPN. As expected, Ebf1 was absent from the entire striatal mantle at E17.5 in both cKOs (Figs. 4b-d) and striatal size was slightly reduced, as in *Ebf1^−/−^* mice^42^. However, all the generic early SPN markers we examined, including Ctip2 (Figs. 4e-g), Foxp1, gad1 (not shown), DARPP-32 (Figs. 8j-l) indicated that the striatum was still formed by GABAergic neurons with a SPN-like identity. In order to compare the phenotypes in the two cKOs and gain insights in the molecular programs controlled by *Ebf1*, we performed RNA-sequencing (RNA-seq) in striatal tissue of control, *Islet1^Cre^;Ebf1^fl/−^* and *Dlx5/6::Cre;Ebf1^fl/−^* embryos at E17.5 (Figs. 4h-k and Tables S1, S2). Importantly, most genes were commonly deregulated in the two cKOs (Figs. 4h-k and Tables S1, S2) suggesting that inactivation of *Ebf1* in dSPN or all SPN lead to similar phenotypes. To compare the datasets, we examined genes that were previously identified as representative of the core SPN generic identity, versus specific of dSPN or iSPN identity^41, 43, 45, 46, 69^. We found that genes associated with a generic SPN identity or with iSPN identity were not significantly altered in both mutants (Figs. 4h-i and Tables S1, S2). In contrast, the expression of genes associated with dSPN were either preserved (*Drd1, Foxp2, Islet1, Pdyn*), or severely deregulated in both mutant models (Figs. 4j-k and Tables S1, S2). The preserved expression of Foxp2 and Islet1 shows that these transcriptional regulators must act in a partially distinct pathway. Deregulated genes encoded selective transcriptional regulators such as *Zfp521, Mef2c*^31, 44, 70–72^, transmembrane receptors such as *PlexinD1*, which is involved in synaptogenesis^27^, and intracellular effectors such as *Slc35d3*, which regulates D1 receptor trafficking^73^. These results were validated by *in situ* hybridization on selected key genes (Fig. S4). Finally, Gene Ontology analyses revealed specific alterations of factors regulating axon development, cell-cell adhesion properties and synaptogenesis in both cKOs (Figs. S4m-o). Taken together, our findings indicate that, while dispensable for the acquisition of a generic SPN transcriptomic profile, *Ebf1* regulates the expression of genes involved in selective aspects of dSPN differentiation.

**Figure 4.**
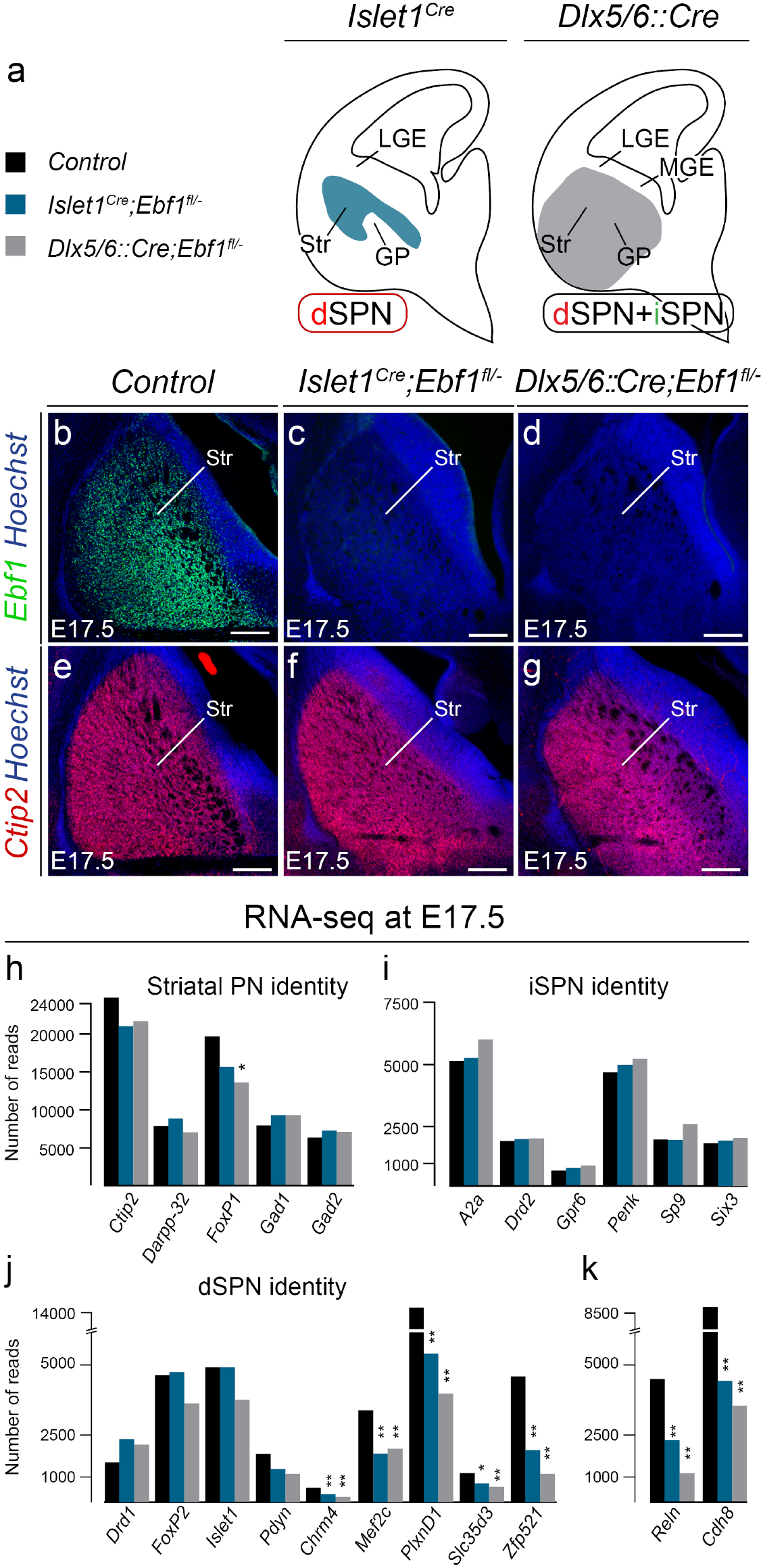
Two *Ebf1* conditional mutants similarly affect dSPN differentiation. (**a**) Schematic representations showing that *Islet1^ŋre^* and *Dlx5/6::Cre* lines drive recombination in differentiated dSPN of the mantle and SVZ/mantle SPN, respectively. (**b-d**) Immunostaining at E17.5 shows a complete absence of Ebf1 protein in the striatum following conditional deletion with either *Islet1^Cre^* or *Dlx5/6::Cre* lines (at least n=3 for each genotype and marker). (**e-g**) Expression of the SPN-specific marker Ctip2 is maintained in both *Ebf1* cKOs. (**h-k**) RNA-sequencing performed at E17.5 on control, *Islet1^Cre^;Ebf1^fl/−^* and *Dlx5/6::Cre;Ebf1^fl/−^* striatal tissues reveals that genes implicated in SPN (h) or iSPN identity (i) are unaffected whereas subsets of genes associated with dSPN identity (j) as well as the striatum-enriched genes *Reln* and *Cdh8* (k) are drastically down-regulated in both cKOs. Importantly, *Foxp2* transcription is not significantly affected by *Ebf1* cKO at E17.5. * p<0.05: **p<0.001. Scale bar equals 200 μm. GP, globus pallidus; LGE, lateral ganglionic eminence; MGE, medial ganglionic eminence; Str, striatum.

### *Ebf1* conditional deletion selectively impairs dSPN differentiation

Since our transcriptomic analysis revealed that *Ebf1* controls the prenatal expression of major regulators of axon development and synaptogenesis (Figs. 4 and S4), we examined striatonigral and striatopallidal projections in both cKOs (Fig. 5). In P5 controls, DARPP-32+ labels SPN axons including the ones of dSPN which crossed the globus pallidus (GP) and the entopeduncular nucleus (EP), joined the cerebral peduncle (CP) and reached the substantia nigra (SN) (Figs. 5a). In both cKOs, DARPP-32+ projections extended to the EP but only a small fraction of them reached the SN (Figs. 5a-c), with axonal density in the CP showing an approximatively 60% reduction compared to controls (Figs. 5d-g). In contrast, when we examined striatopallidal projections by measuring the density of Enkephalin signal in the GP we found no major differences between cKOs and controls (Figs. 5h-k). Thus, *Ebf1* inactivation selectively impaired the capacity of dSPN to form the direct pathway.

**Figure 5.**
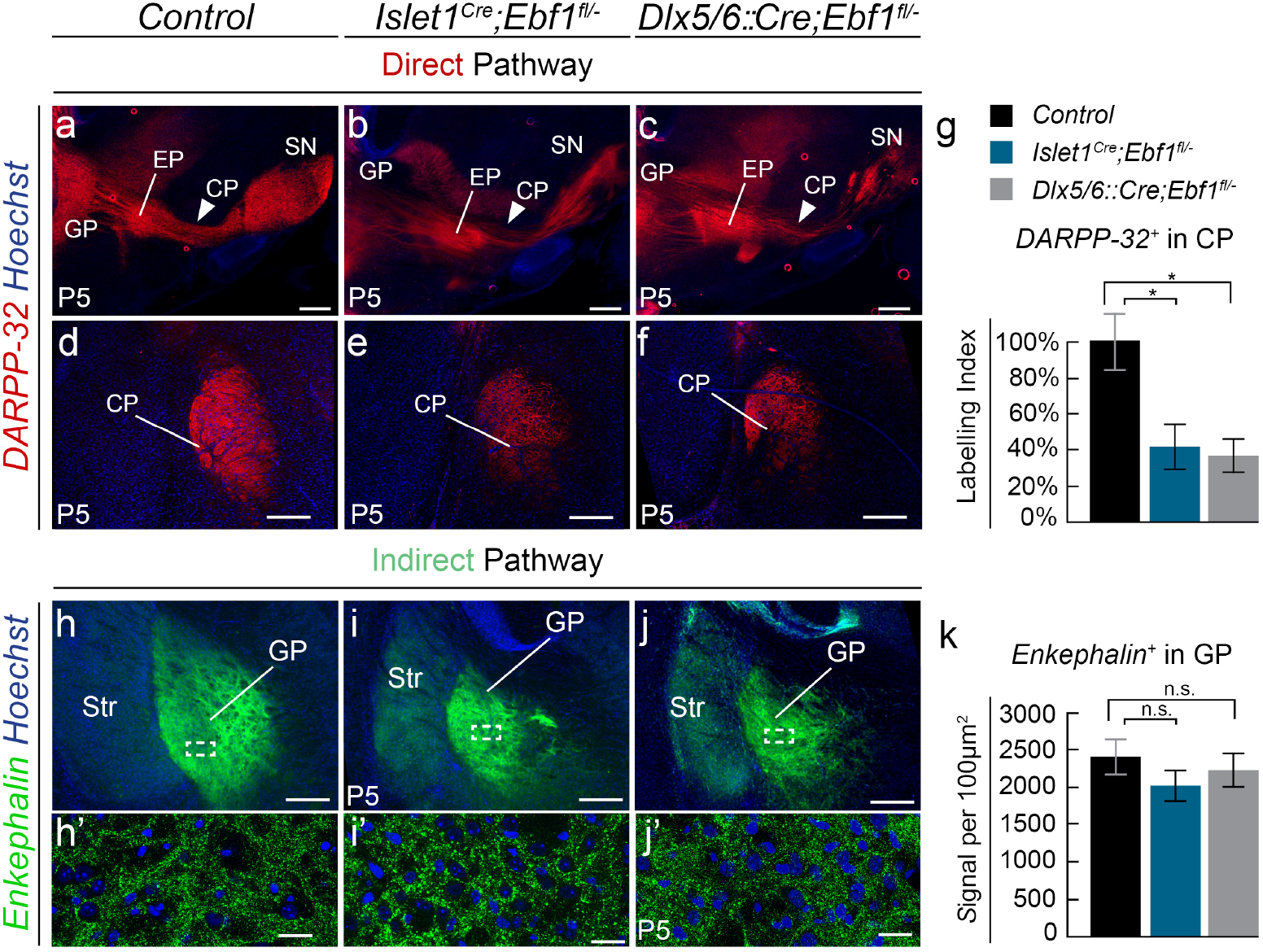
*Ebf1* conditional inactivation affects direct pathway formation. (**a-c**) DARPP-32 staining of striatal axons on P5 sagittal sections reveal that, instead of reaching their target in controls (n=3) (a), striatofugal axons show limited extension passed the entopeduncular nucleus (EP) in both *Islet1^Cre^;Ebf1^f?/−^* (n=4) (b) and *Dlx5/6::Cre;Ebf1^fl/−^* (n=4) (c) mutants (solid arrowheads). (**d-g**) DARPP-32 staining in coronal sections showing reduced axonal projections in the cerebral peduncle (CP) of *Ebf1* cKOs, quantified in (g). Labelling Index is a function of signal area and mean intensity (p=0.028 for *Islet1^Cre^;Ebf1^fl/−^* and p=0.023 for *Dlx5/6::Cre;Ebf1^fl/−^).* (**h-k**) Indirect pathway terminals in GP, labelled by Enkephalin immunostaining show similar densities in controls and both *Ebf1* cKOs, as quantified in (k) (n=3 mice for each condition, p>0.05 for controls versus both *Islet1^Cre^;Ebf1^fl/−^* and *Dlx5/6::Cre;Ebf1^f/−^).* Results are presented as mean values ± s.e.m. Scale bars equal 400 μm (a-c), 200 μm (d-j) and 25 μm (h’-j’). CP, cerebral peduncle; EP, enteropeduncular nucleus; GP, globus pallidus; SN, substancia nigra; Str, striatum.

In contrast to Ebf1^−/−^^42–44^, cKOs survive up to adult age, allowing us to examine the long-term impact on SPN properties and pathway formation. We observed in cKOs that dtTomato-labeled dSPN were reduced in number and size at P45 in cKO mice (Figs. 6a-g), indicating possible functional impairment. To further analyze the impact of *Ebf1* inactivation on dSPN connectivity, we recorded glutamatergic miniature excitatory postsynaptic currents (mEPSCs) in the wholecell configuration from adult control and cKO mice. We used either dtTomato or *Drd2-EGFP* expression as selective SPN subtype markers (Figs. 6h-j). Moreover, neurons were filled with biocytin during recording for later morphology reconstructions. Consistent with dtTomato labelling, reconstruction of mutant biocytin-filled neurons showed overall reduced dSPN morphology, in contrast to iSPN (Figs. 6k-m). In addition, in both cKOs, dSPN mEPSC frequency was severely reduced, with most cells presenting no detectable spontaneous synaptic activity. In contrast, iSPN did not show significant differences from controls (Figs. 6g-l and Fig. S5a-f). We further examined the functional impact on direct and indirect pathways using a pharmacological approach *in vivo*, by testing motor responses to injections of either D1 or D2 receptor agonists^41, 74^. In baseline conditions, we found that neither cKOs showed major locomotion deficits, *Dlx5/6::Cre; Ebf1^fl/−^* mutants exhibiting only a mild hyperactivity phenotype (Fig. S5g). Such phenotype, which is likely due to extra-striatal defects, precludes a full interpretation of pharmacological challenges (Fig. S5). We thus focused on *Islet1^Cre^; Ebf1^fl/−^* mutants and first activated the direct pathway by injection of the D1 receptor agonist SKF82958 (Fig. 6n). In contrast to control mice, D1 receptor agonist injections in *Ebf1* cKOs did not lead to significant locomotion increases, indicating important functional disruptions of the direct pathway in line with the anatomical defects observed (Figs. 5a-f). Conversely, activation of the indirect pathway through injections of the D2 receptor agonist quinpirole (Fig. 6o) caused strong, statistically similar decreases in the mobility in both control and cKOs, confirming that direct pathway functions are selectively affected by *Ebf1* inactivation. Notably, we found that the indirect pathway is also functionally preserved in *Dlx5/6::Cre; Ebf1^fl/−^* mutants (Fig. S5h).

**Figure 6.**
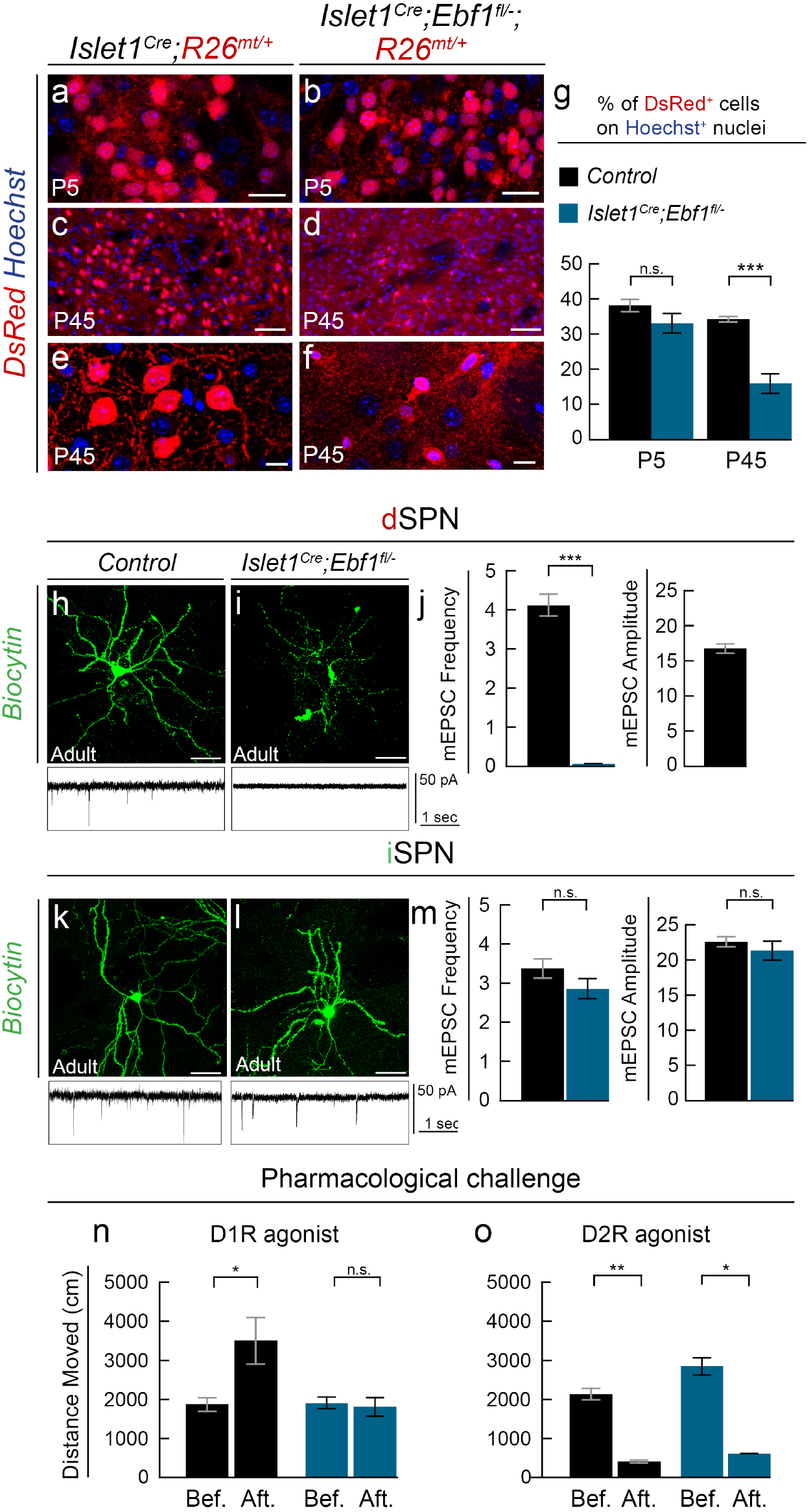
*Ebf1* cKO have specific impairments in dSPN functioning. (**a-g**) tdTomato+ labeling in dSPN cell bodies of *Islet1^Cre^;Ebf1^fl/−^;R26^mt/+^* mice is relatively preserved at early postnatal stages (a-b) but severely altered at later stages (c-f), as quantified in (g); at P5, the percentage of DsRed+ cells on the Hoechst nuclei is 38±2% in Control striata, 33±3% in *Islet1^Cre^;Ebf1^fl/−^* (p>0.05). At P45, Control: 34±1%, *Islet1^Cre^;Ebf1^fl/−^* 16±3% (p=0.002). n=3 mice for each condition. Results are presented as mean values ± s.e.m. (**h-j**) Patch-clamp recordings of dSPN in acute slices obtained from control (33 cells) and *Islet1^Cre^;Ebf1^fl/−^;R26^mt/+^* (20 cells) adult mice highlight a drastic decrease of mEPSC frequency in mutants (sample tracks in g-h, quantification of mEPSC in i). (**k-m**) Conversely, iSPN recorded in controls (27 cells) and *Islet1^Cre^;Ebf1^fl/−^; R26^mt/+^* (14 cells) slices show no relevant differences in mEPSC amplitude and frequency. mEPSC amplitudes could not be quantified in mutant dSPN because of the extremely low number of events recorded. Results are presented as mean values ± s.e.m. (**n-o**) Adult *Islet1^Cre^;Ebf1^fl/−^* mice show no motor response to D1R agonist SKF38393 injections (n), whereas the D2R agonist quinpirole induces depression of locomotion similarly to control mice (o). This is shown by comparing the total distance travelled in 10 minutes in the open field before (Bef.) and 40 minutes after (Aft.) drug injections (n_control_=8 and n_cKO_=7). Results are presented as mean ± standard deviation. Scale bars equal 20 μm (a, b, e, f), 100 μm (c, d) and 40 μm (g-k), respectively. * p-value <0.05, ***p-value <0.0001.

Our results reveal that development and functioning of the direct pathway is selectively impaired in adult *Ebf1* cKOs, while cardinal properties of iSPN, including circuit integration and electrophysiological properties, are preserved. Thus, iSPN differentiation and maturation are largely independent of dSPN.

### Ebf1 deletion affects iSPN migration and intermix in the striatal matrix

Next, we examined whether, in addition to its role in dSPN differentiation, Ebf1 could play a role in SPN intermix and striatal mosaic formation. As is the case for other dSPN markers such as Islet1 and Drd1 (Figs. 4 and S4), the transcriptional levels of FoxP2 were preserved in both early *Ebf1* cKOs (Figs. 4 and S4). Therefore, we used its expression to analyze the distribution of dSPN in the embryonic striatum. We consistently observed non-overlapping populations of Ctip2+ FoxP2+ dSPN and Ctip2+ Drd2-EGFP+ iSPN in mutant striata (Fig. 7). However, we found that dSPN and iSPN distribution was strikingly different in the striatum of controls and of the two cKOs at E17.5 (Fig. 7). Indeed, on one side the density of dSPN in both cKOs was reduced in the dorsal striatum and increased in the lateral region in comparison to controls (Figs. 7a-e). Conversely, the density of Drd2-EGFP+ iSPN was enhanced dorsally and reduced laterally (Figs. 7f-i). Thus, although the overall respective proportions of dSPN and iSPN were preserved, *Ebf1* inactivation perturbed the distribution of the two SPN populations.

**Figure 7.**
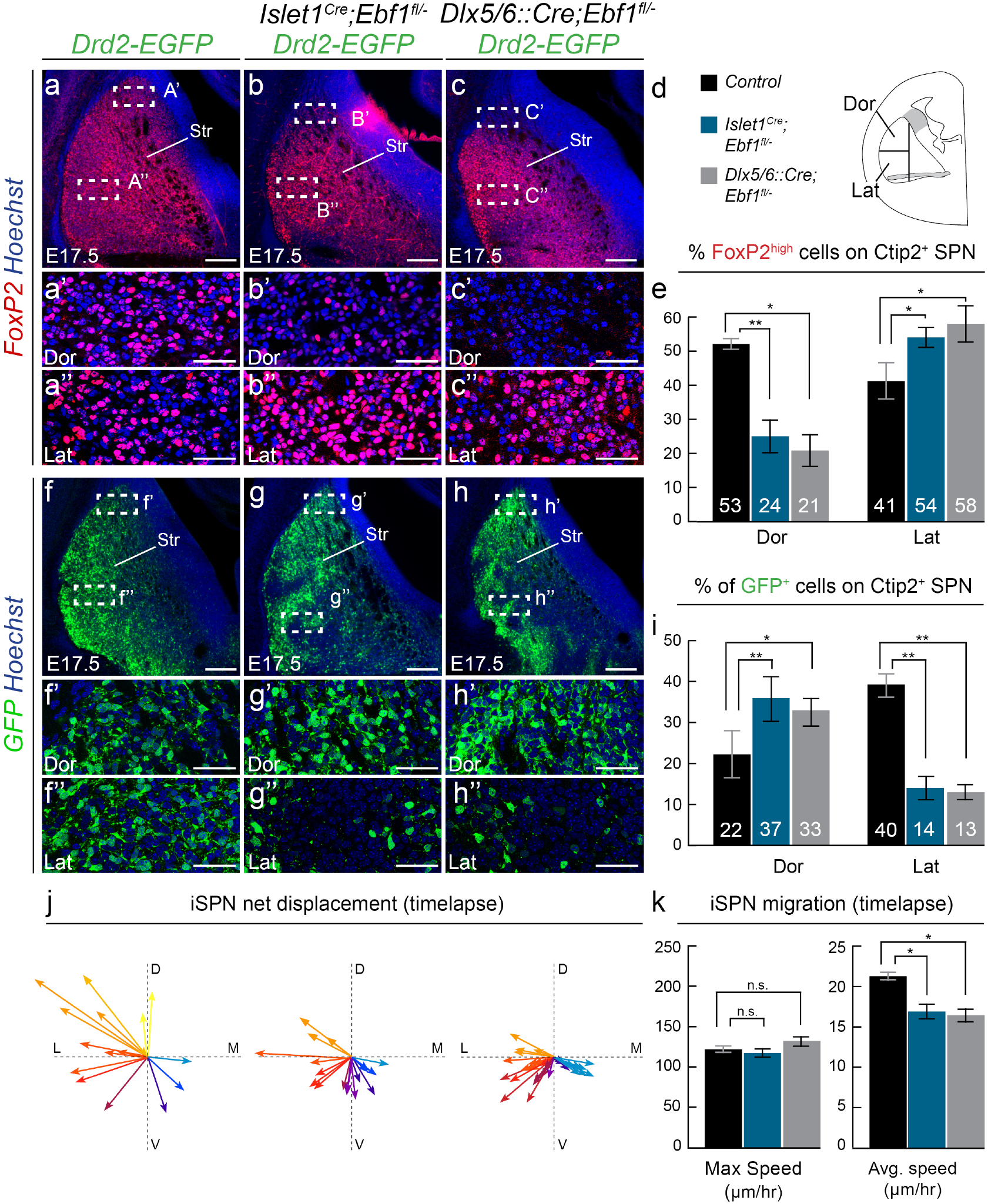
*Ebf1* cKO affects iSPN migration and intermix with dSPN. (**a-e**) Distribution of FoxP2^high^; Ctip2+ dSPN is altered in *Ebf1* cKOs at E17.5, with lower density in the dorsal (Dor) (a’-c’) and higher in the lateral (Lat) part (a”-c”) part of the striatum (Str); the two regions are defined in (d) (n=3 for each condition). (e) In control Dor Str, 53±3.5% of all Ctip2+ SPN are Foxp2+, 24±6% (p =0.0079) in *Islet1^Cre^;Ebf1^fl/−^* and 21±8%. (p=0.03) in *Dlx5/6::Cre;Ebf1^fl/−^.* In control Lat Str, Foxp2+ cells form 41±11% of Ctip2+ SPN, raising to 54±3% in *Islet1^Cre^;Ebf1^fl/−^* (p=0.01) and to 58±11% in *Dlx5/6::Cre;Ebf1^fl/−^* (p=0.03). (**f-h”**) Distribution of Drd2-EGFP+;Ctip2+ iSPN is altered by *Ebf1* cKO, with higher density in Dor Str (f’-h’) and lower in Lat Str (f”-h”). (i) Quantification of iSPN distribution. In control Dor Str, 22±12% of Ctip2+ SPN are GFP+. This percentage increases to 37±11% in *Islet1^Cre^;Ebf1^fl/−^;Drd2-EGFP* (p=0.007) and to 33±7% in *Dlx5/6::Cre;Ebf1^fl/−^;Drd2-EGFP* (p=0.03). Conversely, in the Lat Str, the percentage of of double-positive GFP^+^Ctip2^+^ iSPN equals 40±6% in controls, 14±6% in *Islet1^Cre^;Ebf1^fl/−^;Drd2-EGFP* (p=0.008) and 13±4% in *Dlx5/6::Cre;Ebf1^fl/−^; Drd2-EGFP* embryos (p<0.008). (**j-k**) Analysis of iSPN migration at E15.5 in acute slices of *Islet1^Cre^;Ebf1^fl/−^; Drd2-EGFP* and *Dlx5/6::Cre;Ebf1^fl/−^;Drd2-EGFP* embryos. (j) Multidirectional displacement vectors from sample video tracking of control, *Islet1^Cre^;Ebf1^fl/−^; Drd2-EGFP* and *Dlx5/6::Cre;Ebf1^fl/−^;Drd2-EGFP* iSPN (colors illustrate directionality). (k) Trajectory analysis (*Islet1^Cre^;Ebf1^fl/−^; Drd2-EGFP:* 58 cells tracked, n=3 independent experiments; *Dlx5/6::Cre;Ebf1^fl/−^;Drd2-EGFP* 29 cells tracked, n=2 independent experiments) shows reduction of iSPN average speed after 180’ in *Ebf1* cKOs (17,25±1 μm/hr for *Islet1^Cre^;Ebf1^fl/−^; Drd2-EGFP*, p=0.02; 16,56±1 μm/hr for *Dlx5/6::Cre;Ebf1^fl/−^;Drd2-EGFP* p=0.03) without significant alteration of maximum speed (117,8±5 μm/hr and 131±6 μm/hr, respectively; p>0.05 for both) when compared to control iSPN. Results are presented as mean ± standard deviation (e, i) or s.e.m (k). Scale bars equal 200 μm (a-c, f-h), 50 μm (a’-c” and f’-h”). Dor, Dorsal; Lat, lateral; Str, striatum.

The dorsal accumulation of iSPN near progenitor zones raised the possibility that their migration might be indirectly impaired in *Ebf1* cKO. To test this hypothesis, we first performed timed EdU injections to follow SPN progression into the striatal mantle of *Ebf1* mutants, using Drd2-EGFP and either Foxp2 or dtTomato as respective markers of iSPN and dSPN (Fig. S6 and data not shown). We observed that EdU-stained iSPN abnormally accumulated in the dorsolateral part of the striatum in cKOs (Fig. S6), suggesting that Ebf1 non cell-autonomously regulates their migration. To directly monitor iSPN migration, we performed two-photon time-lapse imaging in E15.5 *Islet1^Cre^;Ebf1^fl/−^;Drd2-EGFP* and *Dlx5/6::Cre;Ebf1^fl/−^;Drd2-EGFP* slices. We found that in both genetic backgrounds, mutant iSPN still harbored pattern of multidirectional migration as in controls (Fig. 7j and Movies S2-4). However, analyses over several time-lapse movies showed that iSPN average speed was significantly decreased (Fig. 7k), thereby revealing that Ebf1 non cell-autonomously controls the migration efficiency of iSPN. Thus *Ebf1* inactivation in dSPN impairs iSPN migration and their process of intermixing. Since Ebf1 and Islet1 both control dSPN differentiation, we investigated the specificity of such intermixing defect by comparing the distribution of D2 receptor (Drd2) transcripts in *Ebf1* and *Islet1* cKOs (Fig. S7). In contrast to *Ebf1* cKO, we found that iSPN distribution was not drastically altered in *Islet1* cKO (Fig. S7). Therefore, iSPN intermixing specifically relies on Ebf1 expression in dSPN, indicating a novel and specific role for this transcription factor in the acquisition of dSPN properties.

In addition to changes in dorso-ventral distribution, we observed that iSPN presented a higher degree of segregation, especially in the lateral region (Figs. 7f-h”). This observation raised questions about possible disruptions of the striosomes/matrix organization. Using early striosome markers such as DARPP-32^9, 75^ and Fig. S1), we found that the two compartments could be unambiguously identified in both cKOs (Figs. 8a-c). However, we observed that the matrix contained Enkephalin^high^ domains (Figs. 8d-f) which segregated from cellular islands containing matrix dSPN (Fig. S8) in both cKOs, suggesting an intermixing deficit between iSPN and dSPN. To further test this possibility, we examined Drd2-EGFP controls and cKOs at E17.5, when compartment segregation is largely engaged^20, 21, 47–51^ and proto-striosomes can be identified by the expression of SubstanceP (not shown) and DARPP-32 (Figs. 8j-l), a maturation marker, which is turned on in the matrix only at later stages. We observed that the deficit in iSPN and dSPN intermixing was more pronounced in the matrix (Figs. 8k-l’), consistently with previous reports showing that *Ebf1* inactivation more specifically affects matrix dSPN^44^. Collectively, our findings reveal that the intermixing of iSPN and dSPN in the matrix specifically relies on Ebf1 expression in dSPN (Fig. 8m-n). This major step in the emergence of a balanced striatum parallels the formation of compartments, thereby revealing that multiple migratory processes regulate the assembly of the striatal mosaic.

**Figure 8.**
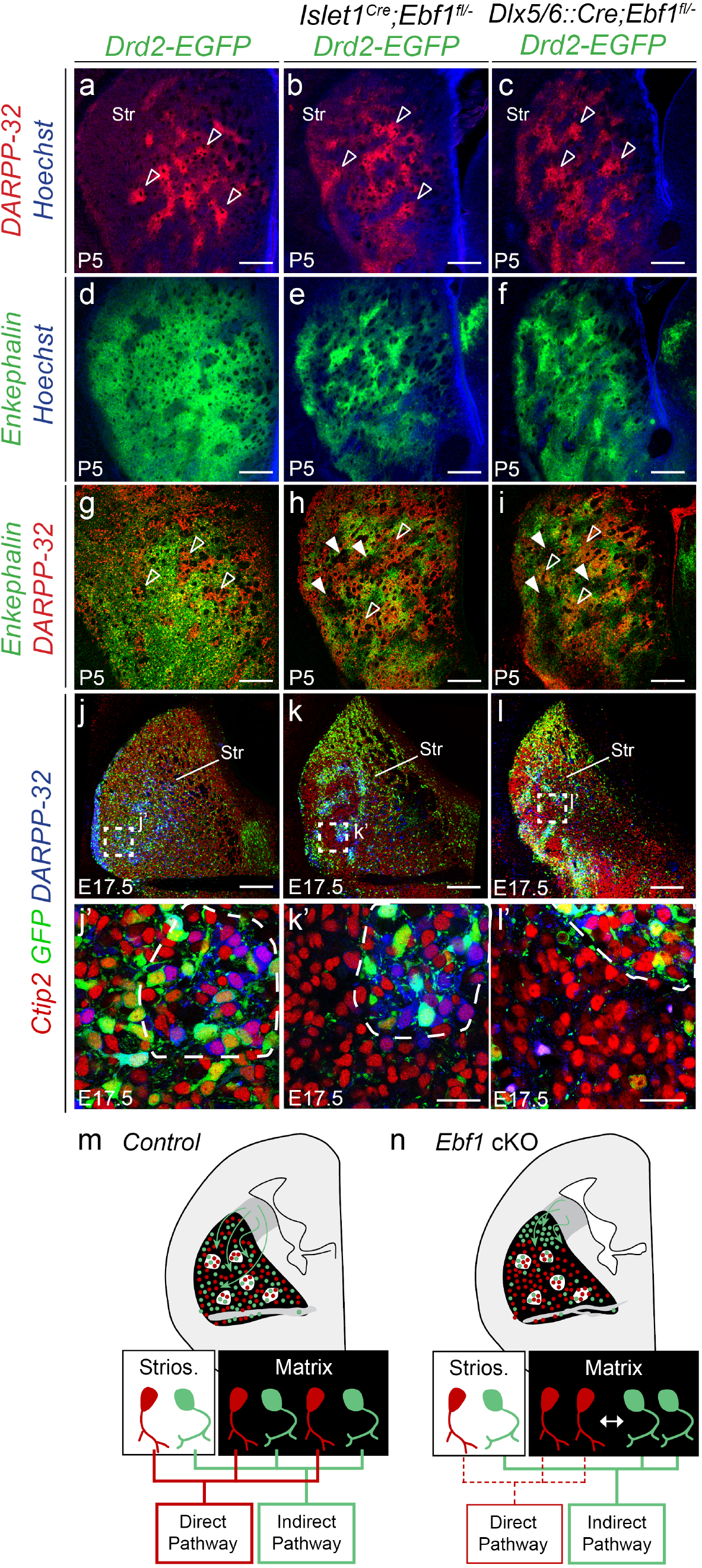
Defective intermixing is observed in the matrix compartment. (**a-c**) DARPP-32 staining in P5 striatal coronal sections reveals that striosomes are still recognizable in *Islet1^Cre^;Ebf1^fl/−^* and *Dlx5/6::Cre;Ebf1^fl/−^* (empty arrowheads) (n=3 for each genotypes and markers). (**d-i**) Conversely, in *Ebf1* cKOs the matrix compartment is parceled in Enkephalin-rich and Enkephalin-poor (full arrowheads) areas. (**j-l**’) Intermixing anomalies are prominent in the matrix compartment of both cKOs. In the lateral striatum, iSPN are intermixed both within and outside DARPP-32+ striosomes in controls (j’), whereas iSPN are intermixed in striosomes (delineated by dotted lines) but not in the matrix of *Ebf1* cKOs (j’-k ‘). (**m-n**) Schematic representation of the functions of Ebf1 in dSPN differentiation and in the non-cell autonomous regulation of iSPN migration. n=3 for each genotype. Scale bars equal 200 μm (a-l) and 30 μm (j’-l’). Str, striatum.

## Discussion

Striatal functioning depends on the balanced activation of intermixed direct and indirect pathway neurons^76^. It is thus essential to understand how the two subtypes are specified and how they assemble into a mosaic striatal architecture. Here, we show that while dSPN and iSPN differentiation is largely independent, their intermixing relies on a developmental crosstalk, which is crucial to organize striatal architecture and is largely independent from striosome and matrix compartment formation.

While both SPN derive from the lgE^19, 22–24, 49^, producing fully differentiated dSPN or iSPN *in vitro*^77–81^ has remained a challenge. While there was increasing evidence that distinct transcriptional programs underlie the specification of both subtypes^23, 41, 45, 46^, *in vitro* studies raised questions regarding the mechanisms controlling SPN differentiation. Our results confirm that dSPN and iSPN possess distinct genetic fingerprints from early embryonic stages, reinforcing the idea that the two cell types are already specified when they enter the striatal mantle.

Furthermore, iSPN progressively invade dSPN-rich areas through dynamic multidirectional migration. Intriguingly, iSPN are initially detected in the lateralmost part of the striatum, consistently with the observation that transcription factors regulating iSPN development, Six3 and Sp9, are located in the dorsal part of the LGE^23^. Tangential migration is a well-known mechanism for increasing neuronal diversity in different brain structures. In the telencephalon, tangential migration has mostly been associated with interneuron behavior^82, 83^, but has also been involved in the positioning of other LGE-derived projection neurons, including arkypallidal neurons of the globus pallidus^84, 85^, corridor neurons^86–88^ and dSPN-like neurons of the central^89^ and extended amygdala^90^. Further investigation will be required to determine the cellular and molecular mechanisms governing the migration and interaction of different LGE-generated populations. Nevertheless, tangential migration of projection neurons is emerging as a shared mechanism to build and increase cellular diversity in basal ganglia nuclei, in contrast to the laminated cerebral cortex where projection neurons migrate radially from the underlying proliferative neuroepithelium^91^.

Regarding the transcriptional cascades governing SPN identity, our work indicates that Ebf1 plays a major role in dSPN differentiation, independently of the other known regulators Islet1 and Foxp2. *Ebf1* is a member of the COE family that plays a key role in the specification of B cells and the differentiation of neuronal subtypes^79, 92–100^. Ebf1 acts both as a direct regulator of downstream target genes^92, 97, 98, 100^ and as a chromatin remodeler, poising or regulating the accessibility of enhancers^101, 102^. *Ebf1* is expressed in the developing striatum^42^ and analyses of *Ebf1* full knockout mice showed normal proliferation in the LGE but defective survival of matrix dSPN and disorganized projections to the SN^43, 44^. By combining analysis of two distinct cKOs, we first showed that Ebf1 is dispensable for the fate choice between dSPN and iSPN identity and the core aspects of differentiation into GABAergic SPN neurons. Indeed, *Ebf1* inactivation leads to an abnormal differentiation of dSPN-like neurons that nonetheless retain their GABAergic identity and expression of DARPP-32, Ctip2, FoxP1, Drd1, Islet1 and FoxP2. Notably, deregulated genes included *Zfp521* and *Mef2c*, which are directly bound by Ebf1 in pro-B cells and thus likely direct targets^92, 97^. Second, *Ebf1* is not expressed in the ventral striatum and its inactivation selectively perturbs the matrix compartment of the dorsal striatum, potentially revealing a subdivision of dSPN neurons. Third, *Ebf1* regulates expression of genes controlling axon and synaptogenesis, thereby highlighting a major role in circuit integration. Importantly, *Ebf1* and *Islet1* both regulate direct pathway formation and the expression of *PlexinD1*^27, 41, 42, 45^, suggesting that the two genes might cooperatively regulate specific features of dSPN neurons.

Finally, we found that *Ebf1* non cell-autonomously regulates iSPN intermixing with dSPN in the matrix compartment. In *Ebf1* cKOs, iSPN are correctly specified and functional, but their progression and intermixing with dSPN is impaired. This intermixing deficit occurs independently of the formation of striosomes/matrix and is more pronounced in the matrix compartment. Remarkably, this phenotype is absent in *Islet1* cKO, indicating a novel and specific role for Ebf1 in the building of striatal functional organization. The underlying mechanisms remain to be deciphered and could include the production of membrane-bound or secreted factors acting on iSPN. Our study also reveals that the intermix of dSPN/iSPN and the formation of striosomes/matrix compartments constitute parallel processes. Indeed, it has been shown that early-born and late-born SPN interact in late embryogenesis to respectively form the striosomes and the matrix via migration and ephrinA/EphA4-dependant cell-sorting^15, 20, 21, 24, 47, 49–51, 103^. In *Ebf1* cKOs, striosomes do form but the intermix of matrix neurons is defective. Our study thus reveals that distinct steps of migration and reorganization control the emergence of two compartments comprising intermixed dSPN/ iSPN (Fig. S9), an architecture which is essential for striatal functioning.

Taken together, our work shows that while iSPN and dSPN specification is largely independent, the assembly of striatal mosaic emerges from a dSPN-dependent tangential migration of iSPN. This study establishes a novel framework for the formation of the striatum, a major structure associated with developmental disorders, and provides key insights on how migration controls the wiring of neural circuits.

## Methods

### Mouse lines

For fate mapping studies, *Islet1^Cre/+^* animals^63^ were crossed with either *R26^mt/mt^* or *R26^mt/mt^;Drd2-EGFP* mice^52^. To obtain *Islet1^Cre/+;^Ebf1^fl/−^, Islet1^Cre/+^;Ebf1^fl/−^ R26^mt/+^* and *Islet1^Cre/+^:Ebf1^fl/−^;Drd2-EGFP* conditional mutants, we initially crossed *Islet1^Cre/+^* mice with *Ebf1^+/−^*^104^ to generate *Islet1^Cre/+^;Ebf1^fl/−^* animals. These were in turn backcrossed with either *Ebf1^fl/fl^*^97, 105^, *Ebf1^fl/fl^;R26^mt/mt^*, or *Ebf1^fl/fl^;Drd2-EGFP* mice, respectively. Similarly, *Dlx5/6::Cre;Eb1^fl/−^* and *Dlx5/6::Cre;Ebf1^fl/−^;Drd2-EGFP* mice were obtained by crossing *Dlx5/6::Cre*^65^ mice with *Ebf1^+/−^* to generate *Dlx5/6::Cre;Ebf^+/−^* animals, which were in turn backcrossed with *Ebf1^fl/fl^* or *Ebf1^fl/f^;Drd2-EGFP* mice. All transgenic lines were maintained on a C57/Bl6 background, with the exception of *Islet1^Cre/+^* and *Drd2-EGFP* lines that remained on a B6D2F1/J genetic background. Heterozygous embryos did not show any phenotype and were used as controls. *Nkx2.1^Cre/+^* mice provided by S. Anderson’s laboratory^106^. 12-μm-thick cryosections from *Dlx1^Cre/+^;Islet1^fl/fl^* embryos provided by K. Campbell’s laboratory^41^. The day of vaginal plug was considered as embryonic day (E) 0.5 and day of birth as postnatal day (P) 0. Animals were bred under French and EU regulations, following recommendations of the local ethics committee.

### *In situ* hybridization and immunohistochemistry

For *in situ* hybridization, brains were fixed overnight in 4% paraformaldehyde in PBS (PFA) at 4°C. 80 to 100μm-thick free-floating vibratome sections (Leica S1000) were hybridized as described^105^; *Dlx1^Cre/+^;Islet1^fl/fl^* and respective control cryosections were hybridized as described^107^. For immunohistochemistry, mice were perfused with 4% PFA. Brains were dissected and post fixed overnight at 4°C. Immunohistochemistry was performed on 60μm-thick free-floating vibratome sections. Slices were incubated 1h at room temperature (RT) in a blocking solution containing 0,25% Triton X-100 (Sigma), 0,02% Gelatine in PBS, and incubated in the same blocking solution with primary antibodies overnight at 4°C. Primary antibodies were used at the following concentrations: rat anti-CTIP2 1/500 (Abcam), mouse anti-DARPP-32 1/100 (Santa Cruz), rabbit anti-DARPP-32 1/1000 (Millipore), rabbit anti-DsRed 1/500 (Living colors), rabbit anti-Enkephalin 1/500 (Millipore), rabbit anti-Ebf1 1/250 (Abcam), rabbit anti-FoxP1 1/200 (Abcam), goat anti-FoxP2 1/200 (Santa Cruz), chicken anti-GFP 1/1000 (Aves), rabbit anti-Slc35d3 1/250 (Novusbio), rat anti-SubstanceP 1/400 (Millipore), rabbit anti-Tyrosine Hydroxylase 1/1000 (Abcam), guinea pig anti-vGlut1 1/10000 (Millipore), guinea pig anti-vGlut2 1/10000 (Millipore). Sections were rinsed several times in PBS and incubated from 2h to overnight at 4°C with the appropriate fluorescent secondary antibodies: A488-conjugated donkey antirabbit, anti-rat or anti-chicken, Cy3-conjugated donkey anti-rat, anti-goat, antimouse, Cy5-conjugated donkey anti-rabbit or anti-rat, DyLight488-conjugated donkey anti-guinea pig (1/400, all antibodies from Jackson ImmunoResearch). Hoechst (Sigma) was used for fluorescent nuclear counterstaining and sections were mounted in Mowiol or Vectashield (Vector).

### Birthdating

Pregnant dams were injected intraperitoneally at the appropriate gestation day with a solution containing 5-Ethynyl-2’-deoxyuridine (EdU, Thermo Fisher). 60-100μm-thick free-floating vibratome sections were processed following manufacturer instructions (Click-iT EdU Alexa Fluor 488 Imaging kit, Life Technologies) for 30 minutes at RT. Sections were rinsed three times in 3% BSA and then in PBS. Hoechst staining was performed for 30 min at RT before pursuing the immunohistochemistry protocol as described above.

### Image acquisition, analysis and quantification

Images were acquired on a fluorescence microscope (Leica MZ16 F), a fluorescent microscope (Leica DM5000 B) or a confocal microscope (Leica TCS SP5). Images were then processed with ImageJ and Adobe Photoshop software. For cell density and colocalization analysis, three different rostro-caudal striatal levels of the striatum were initially defined, using the anterior commissure as an anatomical landmark. Single-plane confocal images were taken at each level in three different animals for each condition. For cell distribution analysis in Fig. 7, a high-definition tilescan of the whole striatum at medial level was imaged on a single confocal plane in at least three different animals for each condition. The tilescans were later subdivided in lateral and dorsal regions (Fig. 7d) for quantification. Cell counting was performed semi-automatically using built-in functions in ImageJ. For each experiment, sample images were manually counted to double-check the quality of semi-automated counting.

### Timelapse imaging and analysis

E13.5 and E15.5 *Drd2-EGFP* and *Dlx5/6::Cre;Ebf1^fl/−^;Drd2-EGFP* embryos were dissected and kept in an ice-cold solution consisting of L-15 Medium (SIGMA-Aldrich) supplemented with 3% Glucose. Brains were included in 3,5% low-melting agarose (Promega) in L-15 Glucose+ solution and 350 μm-thick coronal slices were cut on a vibratome. Slices were imaged using a multi-photon microscope (DIMM, IBENS Imaging Platform) over a period of up to 240’, while being constantly perfused with a L-15 Glucose+ solution at 37°C and bubbled with O_2_/CO_2_ (95% / 5%). 100 μm-thick z-stacks were imaged every 6’ for each single frame. For time-lapse movie analysis, eventual drift in the three dimensions was first corrected using IMARIS software (Bitplane, Oxford); subsequently, z-stacks were flattened on a two-dimensional plane. The high number of cells present in each single movie rendered automated analysis of cell displacement unadvisable. Therefore, single cell displacement was measured manually frame-by-frame using the “Manual Tracking” plugin in ImageJ. Cells included in the analysis responded to two criteria: being readily identifiable from the first to the last frame of the movie and showing a total net displacement of more than 50 μm. In order to examine directionality and migratory pattern of iSPN migration, stationary (total net displacement less than 50 μm) cells were not included in the analysis.

### RNA-sequencing

E17.5 *Dlx5/6::Cre;Ebf1^fl/−^, Islet1^Cre/+^;Ebf1^fl/−^* and control embryos were dissected in RNAse-free conditions on ice. Brains were conserved in RNAlater stabilization reagent (Qiagen) solution. Following genotype identification via PCR, messenger RNA was obtained from n=3 brains from each condition using RNeasy mini kit (Qiagen). Library preparation and Illumina sequencing were performed at the Ecole Normale Supérieure Genomic Platform (Paris, France). Messenger (polyA+) RNAs were purified from 400ng of total RNA using oligo(dT). Libraries were prepared using the strand specific RNA-Seq library preparation TruSeq Stranded mRNA kit (Illumina). Libraries were multiplexed by 9 on run Nextseq 500. A 75bp read sequencing was performed on a Nextseq 500 device (Illumina). A mean of 37,5±8,95 million passing Illumina quality filter reads was obtained for each of the 9 samples. Analyses were performed using the Eoulsan pipeline, including read filtering, mapping, alignment filtering, read quantification, normalisation and differential analysis. Before mapping, poly N read tails were trimmed, reads ≤40 bases were removed, and reads with quality mean ≤30 were discarded. Reads were then aligned against the Mus musculus genome from Ensembl version 81 using STAR (version 2.4.0k). Alignments from reads matching more than once on the reference genome were removed using Java version of samtools. To compute gene expression, Mus musculus GFF3 genome annotation version 81 from Ensembl database was used. All overlapping regions between alignments and referenced genes were counted using HTSeq-count 0.5.3. Sample counts were normalized using DESeq 1.8.3. Statistical treatments and differential analyses were also performed using DESeq 1.8.3^108^.

### Slice preparation and electrophysiological recordings

*In vitro* electrophysiological recordings were performed in coronal slices from the dorsal striatum of control animals (either *Islet1^Cre/+^;R26^mt/+^* or *Drd2-EGFP* mice), *Dlx5/6::Cre;Ebf1^fl/−^;Drd2-EGFP* and *Islet1^Cre/+^;Ebf1^fl/−^;R26^mt/+^* mice. Mice were anesthetized with isofluorane before decapitation. After isolation, the portion of the brain containing the striatum was placed in bicarbonate-buffered saline (BBS) at 2–5°C for a few minutes. Slices (300 μm) were then cut using a 7000smz-2 vibratome (Campden Instruments Ltd.). The slicing procedure was performed in an ice-cold solution containing (in mM): 130 potassium gluconate, 15 KCl, 0.05 EGTA, 20 Hepes, 25 glucose, 1 CaCl_2_ and 6 MgCl_2_. Slices were then briefly transferred to a solution containing (in mM): 225 D-mannitol, 2.5 KCl, 1.25 NaH_2_PO_4_, 25 NaHCO_3_, 25 glucose, 1 CaCl_2_ and 6 MgCl_2_, and finally stored for the rest of the experimental day at 33 °C in oxygenated BBS, containing: 115 NaCl, 2.5 KCl, 1.6 CaCl_2_, 1.5 MgCb, 1.25 NaH_2_PO_4_, 26 NaHCO_3_ and 30 glucose (pH 7.4 after equilibration with 95% O_2_ and 5% CO_2_). For all recordings, slices were continuously superfused with oxygenated BBS, supplemented with the GABAa receptor blocker SR95531 (Gabazine; 2μM) and with Tetrodotoxin (TTX; 500nM), at 32–34°C. Electrophysiological recordings were performed from either dSPN-dTomato positive, iSPN-dTomato negative, dSPN-GFP negative or iSPN-GFP positive dorsal striatal cells. Cells were patched in the transmitted deep red light with which slices were visualized using a CoolSnap HQ CCD camera (Photometrics) run by Metamorph software (Universal Imaging) and mounted on either a Slicescope (Scientifica), or a BX51 (Olympus) microscope. Before patching, Tomato or GFP positive/negative cells were identified by the presence/absence of somatic fluorescence using LEDs of the corresponding excitation wavelengths (Thorlabs) coupled to the slice chamber via the epifluorescence pathway of the microscope.

### Pharmacology and Behavior analyses

The motor responses of *Dlx5/6::Cre;Ebf1^fl/−^* and *Islet1^Cre/+^;Ebf1^fl/−^* mice and their corresponding controls were examined in a round open field arena (diameter: 38 cm) following injection of pharmacological agents. Minimum two weeks separated tests performed with different drugs on the same cohort of mice. For each condition, animals were acclimated to the experimental luminosity conditions (27–35 lux) of the test room for 1 hr. Following adaptation, baseline motor activity in the open field of each mouse was then measured for 8 minutes before subcutaneous injection with one of the following solutions: 0,9% NaCl (control injection); 2mg/Kg D1 receptor agonist SKF82958 (Sigma); and 1mg/Kg D2 receptor agonist Quinpirole (Sigma). Drugs were administered in a volume of 10 ml/Kg of body weight. Following the injection, animals were left in their cage for 40 minutes. They were then re-introduced into the arena for recording their motor activity over another 8 min-long timeperiod. Post-hoc analysis of the total distance travelled was performed using Ethovision software (Noldus).

### Statistical analyses

All data are presented as mean ± SD or SEM (detailed in each figure legend). Two-tailed non-parametric Mann-Whitney U test was used to compare two distributions in all experiments, with the exception of RNA-seq analysis (see above). All statistical analyses were performed using GraphPad Prism software. P-values are as follows: * p<0.05, **p<0.01, ***p<0.0001.

## Additional information

Supplementary information includes 9 supplementary figures, 4 movies and 2 Tables/Spreadsheets.

## Author contributions

AT and SG designed the study and wrote the manuscript. MAD performed and analyzed electrophysiological recordings. MAD, FM, DP and AT performed motor behavior experiments and analyses. FC and SL performed the RNAseq and transcriptomic analyses. BM provided expertise for two-photon imaging. AT, LL, CM and MK performed all other experiments and analyzed data. IG, PMS, KC and RG provided mouse models. All the authors edited the manuscript. SG supervised the project.

## Acknowledgements

We thank members of the S.G. lab, M. Casado, J.A. Girault, C. Goridis, D. Hervé, C. Lena and E. Valjent for stimulating discussions. We are grateful to C. Auger, D. Souchet, A. Boudjouher and C. Le Moal for excellent technical assistance. We thank E. Thierion and L. Danglot for scientific input. We are grateful to the IBENS Imaging Facility (France BioImaging, supported by ANR-10-INBS-04, ANR-10-LABX-54 MEMO LIFE and ANR-11-IDEX-000-02 PSL* Reseach University, “Investments for the future”; NERF N°2011-45; FRM DGE 20111123023; and FRC Rotary International France), the IBENS Genomics Facility (supported by the France Génomique national infrastructure, funded as part of the “Investissements d’Avenir” program managed by the Agence Nationale de la Recherche, ANR-10-INBS-09) and the IBENS acute transgenesis facility. AT is part of the ENP doctoral program. AT was supported by a doctoral fellowship from Boehringer Ingelheim Fonds (BIF), the ENP program and the Labex Memolife « Investements for the future » (ANR-10-LABX-54 MEMO LIFE and ANR-11-IDEX-0001-02 PSL* Research University). This work was supported by grants from INSERM, CNRS, the ANR program (ANR-12-BVS4-0010-01) and the ERC NImO (616080) to S.G, from the ANR program (ANR-12-JSV4-004) to D.P and from NIH (R01 MH090740) to KC.

## References

1. Cui, G. et al. Concurrent activation of striatal direct and indirect pathways during action initiation. Nature 494, 238–242 (2013).

2. Freeze, B. S., Kravitz, A. V., Hammack, N., Berke, J. D. & Kreitzer, A. C. Control of Basal Ganglia Output by Direct and Indirect Pathway Projection Neurons. J. Neurosci. 33, 18531–18539 (2013).

3. Jin, X. & Costa, R. M. Shaping action sequences in basal ganglia circuits. Curr. Opin. Neurobiol. 33, 188–196 (2015).

4. Jin, X., Tecuapetla, F. & Costa, R. M. Basal ganglia subcircuits distinctively encode the parsing and concatenation of action sequences. Nat. Neurosci. 17, 423–430 (2014).

5. Kozorovitskiy, Y., Saunders, A., Johnson, C. A., Lowell, B. B. & Sabatini, B. L. Recurrent network activity drives striatal synaptogenesis. Nature 485, 646–650 (2012).

6. Kravitz, A. V., Tye, L. D. & Kreitzer, A. C. Distinct roles for direct and indirect pathway striatal neurons in reinforcement. Nat. Neurosci. 15, 816–818 (2012).

7. Vicente, A. M., Galvão-Ferreira, P., Tecuapetla, F. & Costa, R. M. Direct and indirect dorsolateral striatum pathways reinforce different action strategies. Curr. Biol. 26, R267–R269 (2016).

8. Gangarossa, G. et al. Spatial distribution of D1R-and D2R-expressing medium-sized spiny neurons differs along the rostro-caudal axis of the mouse dorsal striatum. Front. Neural Circuits 7, 124 (2013).

9. Matamales, M. et al. Striatal Medium-Sized Spiny Neurons: Identification by Nuclear Staining and Study of Neuronal Subpopulations in BAC Transgenic Mice. PLoS ONE 4, e4770 (2009).

10. Yung, K. k. l. & Bolam, J. p. Localization of dopamine D1 and D2 receptors in the rat neostriatum: Synaptic interaction with glutamate-and GABA-containing axonal terminals. Synapse 38, 413–420 (2000).

11. Brimblecombe, K. R. & Cragg, S. J. Substance P Weights Striatal Dopamine Transmission Differently within the Striosome-Matrix Axis. J. Neurosci. 35, 9017–9023 (2015).

12. Burguière, E., Monteiro, P., Mallet, L., Feng, G. & Graybiel, A. M. Striatal circuits, habits, and implications for obsessive–compulsive disorder. Curr. Opin. Neurobiol. 30, 59–65 (2015).

13. Friedman, A. et al. A Corticostriatal Path Targeting Striosomes Controls Decision-Making under Conflict. Cell 161, 1320–1333 (2015).

14. Gerfen, C. R. The neostriatal mosaic: multiple levels of compartmental organization in the basal ganglia. Annu. Rev. Neurosci. 15, 285–320 (1992).

15. Gerfen, C. R., Baimbridge, K. G. & Thibault, J. The neostriatal mosaic: III. Biochemical and developmental dissociation of patch-matrix mesostriatal systems. J. Neurosci. 7, 3935–3944 (1987).

16. Graybiel, A. M. & Grafton, S. T. The Striatum: Where Skills and Habits Meet. Cold Spring Harb. Perspect. Biol. 7, a021691 (2015).

17. Smith, J. B. et al. Genetic-Based Dissection Unveils the Inputs and Outputs of Striatal Patch and Matrix Compartments. Neuron 91, 1069–1084 (2016).

18. Halliday, A. L. & Cepko, C. L. Generation and migration of cells in the developing striatum. Neuron 9, 15–26 (1992).

19. Hamasaki, T., Goto, S., Nishikawa, S. & Ushio, Y. Neuronal cell migration for the developmental formation of the mammalian striatum. Brain Res. Rev. 41, 1–12 (2003).

20. Newman, H., Liu, F.-C. & Graybiel, A. M. Dynamic Ordering of Early Generated Striatal Cells Destined to Form the Striosomal Compartment of the Striatum. J. Comp. Neurol. 523, 943–962 (2015).

21. Song, D. D. & Harlan, R. E. Genesis and migration patterns of neurons forming the patch and matrix compartments of the rat striatum. Brain Res. Dev. Brain Res. 83, 233–245 (1994).

22. Wichterle, H., Turnbull, D. H., Nery, S., Fishell, G. & Alvarez-Buylla, A. In utero fate mapping reveals distinct migratory pathways and fates of neurons born in the mammalian basal forebrain. Development 128, 3759–3771 (2001).

23. Xu, Z. et al. Sp8 and Sp9 coordinately promote D2-type medium spiny neuron production by activating Six3 expression. Dev. Camb. Engl. (2018). doi:10.1242/dev.165456

24. Kelly, S. M. et al. Radial glial lineage progression and differential intermediate progenitor amplification underlie striatal compartments and circuit organization. bioRxiv 244327 (2018). doi:10.1101/244327

25. Arlotta, P., Molyneaux, B. J., Jabaudon, D., Yoshida, Y. & Macklis, J. D. Ctip2 Controls the Differentiation of Medium Spiny Neurons and the Establishment of the Cellular Architecture of the Striatum. J. Neurosci. 28, 622–632 (2008).

26. Bacon, C. et al. Brain-specific Foxp1 deletion impairs neuronal development and causes autistic-like behaviour. Mol. Psychiatry 20, 632–639 (2015).

27. Ding, J. B., Oh, W.-J., Sabatini, B. L. & Gu, C. Semaphorin 3E–Plexin-D1 signaling controls pathway-specific synapse formation in the striatum. Nat. Neurosci. 15, 215–223 (2011).

28. Enard, W. et al. A humanized version of Foxp2 affects cortico-basal ganglia circuits in mice. Cell 137, 961–971 (2009).

29. Garcia-Calero, E., Botella-Lopez, A., Bahamonde, O., Perez-Balaguer, A. & Martinez, S. FoxP2 protein levels regulate cell morphology changes and migration patterns in the vertebrate developing telencephalon. Brain Struct. Funct. (2015). doi:10.1007/s00429-015-1079-7

30. Hébert, J. M. & Fishell, G. The genetics of early telencephalon patterning: some assembly required. Nat. Rev. Neurosci. 9, 678–685 (2008).

31. Ko, H.-A., Chen, S.-Y., Chen, H.-Y., Hao, H.-J. & Liu, F.-C. Cell type-selective expression of the zinc finger-containing gene Nolz-1/Zfp503 in the developing mouse striatum. Neurosci. Lett. 548, 44–49 (2013).

32. Long, J. E. et al. Dlx1&2 and Mash1 Transcription Factors Control Striatal Patterning and Differentiation Through Parallel and Overlapping Pathways.J. Comp. Neurol. 512, 556–572 (2009).

33. Martín-Ibáñez, R. et al. Helios expression coordinates the development of a subset of striatopallidal medium spiny neurons. Development dev.138248 (2017). doi:10.1242/dev.138248

34. Pei, Z. et al. Homeobox genes Gsx1 and Gsx2 differentially regulate telencephalic progenitor maturation. Proc. Natl. Acad. Sci. U. S. A. 108, 1675–1680 (2011).

35. Rataj-Baniowska, M. et al. Retinoic Acid Receptor β Controls Development of Striatonigral Projection Neurons through FGF-Dependent and Meis1-Dependent Mechanisms. J. Neurosci. Off. J. Soc. Neurosci. 35, 14467–14475 (2015).

36. Reimers-Kipping, S., Hevers, W., Pääbo, S. & Enard, W. Humanized Foxp2 specifically affects cortico-basal ganglia circuits. Neuroscience 175, 75–84 (2011).

37. Wang, B., Waclaw, R. R., Allen, Z. J., Guillemot, F. & Campbell, K. Ascl1 is a required downstream effector of Gsx gene function in the embryonic mouse telencephalon. Neural Develop. 4, 5 (2009).

38. Wang, B., Lufkin, T. & Rubenstein, J. L. R. Dlx6 regulates molecular properties of the striatum and central nucleus of the amygdala. J. Comp. Neurol. 519, 2320–2334 (2011).

39. Wang, B. et al. Loss of Gsx1 and Gsx2 function rescues distinct phenotypes in Dlx1/2 mutants. J. Comp. Neurol. 521, 1561–1584 (2013).

40. Yun, K. et al. Modulation of the notch signaling by Mash1 and Dlx1/2 regulates sequential specification and differentiation of progenitor cell types in the subcortical telencephalon. Development 129, 5029–5040 (2002).

41. Ehrman, L. A. et al. The LIM homeobox gene Isl1 is required for the correct development of the striatonigral pathway in the mouse. Proc. Natl. Acad. Sci. 110, E4026–E4035 (2013).

42. Garel, S., Marín, F., Grosschedl, R. & Charnay, P. Ebf1 controls early cell differentiation in the embryonic striatum. Development 126, 5285–5294 (1999).

43. Lobo, M. K., Karsten, S. L., Gray, M., Geschwind, D. H. & Yang, X. W. FACS-array profiling of striatal projection neuron subtypes in juvenile and adult mouse brains. Nat. Neurosci. 9, 443–452 (2006).

44. Lobo, M. K., Yeh, C. & Yang, X. W. Pivotal role of early B-cell factor 1 in development of striatonigral medium spiny neurons in the matrix compartment. J. Neurosci. Res. 86, 2134–2146 (2008).

45. Lu, K.-M., Evans, S. M., Hirano, S. & Liu, F.-C. Dual role for Islet-1 in promoting striatonigral and repressing striatopallidal genetic programs to specify striatonigral cell identity. Proc. Natl. Acad. Sci. 111, E168–E177 (2014).

46. Zhang, Q. et al. The Zinc Finger Transcription Factor Sp9 Is Required for the Development of Striatopallidal Projection Neurons. Cell Rep. 16, 1431–1444 (2016).

47. Fishell, G. & van der Kooy, D. Pattern formation in the striatum: developmental changes in the distribution of striatonigral neurons. J. Neurosci. 7, 1969–1978 (1987).

48. Fishell, G. & van der Kooy, D. Pattern formation in the striatum: neurons with early projections to the substantia nigra survive the cell death period. J. Comp. Neurol. 312, 33–42 (1991).

49. Hagimoto, K., Takami, S., Murakami, F. & Tanabe, Y. Distinct migratory behaviors of striosome and matrix cells underlying the mosaic formation in the developing striatum. J. Comp. Neurol. 525, 794–817 (2017).

50. Krushel, L. A., Fishell, G. & van der Kooy, D. Pattern formation in the mammalian forebrain: striatal patch and matrix neurons intermix prior to compartment formation. Eur. J. Neurosci. 7, 1210–1219 (1995).

51. Passante, L. et al. Temporal regulation of ephrin/Eph signalling is required for the spatial patterning of the mammalian striatum. Development 135, 3281–3290 (2008).

52. Gong, S. et al. A gene expression atlas of the central nervous system based on bacterial artificial chromosomes. Nature 425, 917–925 (2003).

53. Biezonski, D. K., Trifilieff, P., Meszaros, J., Javitch, J. A. & Kellendonk, C. Evidence for Limited D1 and D2 Receptor Co-Expression and Co-Localization Within the Dorsal Striatum of the Neonatal Mouse. J. Comp. Neurol. 523, 1175 (2015).

54. Morello, F. et al. Frizzled3 Controls Axonal Polarity and Intermediate Target Entry during Striatal Pathway Development. J. Neurosci. Off. J. Soc. Neurosci. 35, 14205–14219 (2015).

55. Madisen, L. et al. A robust and high-throughput Cre reporting and characterization system for the whole mouse brain. Nat. Neurosci. 13, 133–140 (2010).

56. Thibault, D., Loustalot, F., Fortin, G. M., Bourque, M.-J. & Trudeau, L.-É. Evaluation of D1 and D2 dopamine receptor segregation in the developing striatum using BAC transgenic mice. PloS One 8, e67219 (2013).

57. Otsu, Y. et al. Optical monitoring of neuronal activity at high frame rate with a digital random-access multiphoton (RAMP) microscope. J. Neurosci. Methods 173, 259–270 (2008).

58. Elshatory, Y. & Gan, L. The LIM-homeobox gene Islet-1 is required for the development of restricted forebrain cholinergic neurons. J. Neurosci. Off. J. Soc. Neurosci. 28, 3291–3297 (2008).

59. Moreno, N., Domínguez, L., Rétaux, S. & González, A. Islet1 as a marker of subdivisions and cell types in the developing forebrain of Xenopus. Neuroscience 154, 1423–1439 (2008).

60. Wang, H. F. & Liu, F. C. Developmental restriction of the LIM homeodomain transcription factor Islet-1 expression to cholinergic neurons in the rat striatum. Neuroscience 103, 999–1016 (2001).

61. Eisenstat, D. D. et al. DLX-1, DLX-2, and DLX-5 expression define distinct stages of basal forebrain differentiation. J. Comp. Neurol. 414, 217–237 (1999).

62. Ruest, L.-B., Hammer, R. E., Yanagisawa, M. & Clouthier, D. E. Dlx5/6-enhancer directed expression of Cre recombinase in the pharyngeal arches and brain. Genes. N. Y. N 2000 37, 188–194 (2003).

63. Srinivas, S. et al. Cre reporter strains produced by targeted insertion of EYFP and ECFP into the ROSA26 locus. BMC Dev. Biol. 1, 4 (2001).

64. Stenman, J., Toresson, H. & Campbell, K. Identification of two distinct progenitor populations in the lateral ganglionic eminence: implications for striatal and olfactory bulb neurogenesis. J. Neurosci. Off. J. Soc. Neurosci. 23, 167–174 (2003).

65. Zerucha, T. et al. A Highly Conserved Enhancer in the Dlx5/Dlx6Intergenic Region is the Site of Cross-Regulatory Interactions betweenDlx Genes in the Embryonic Forebrain. J. Neurosci. 20, 709–721 (2000).

66. Kobayashi, M., Nakano, M., Atobe, Y., Kadota, T. & Funakoshi, K. Islet-1 expression in thoracic spinal motor neurons in prenatal mouse. Int. J. Dev. Neurosci. Off. J. Int. Soc. Dev. Neurosci. 29, 749–756 (2011).

67. Lee, S. et al. STAT3 promotes motor neuron differentiation by collaborating with motor neuron-specific LIM complex. Proc. Natl. Acad. Sci. U. S. A. 110, 11445–11450 (2013).

68. Qu, Q. et al. High-efficiency motor neuron differentiation from human pluripotent stem cells and the function of Islet-1. Nat. Commun. 5, 3449 (2014).

69. Gokce, O. et al. Cellular Taxonomy of the Mouse Striatum as Revealed by Single-Cell RNA-Seq. Cell Rep. 16, 1126–1137 (2016).

70. Chen, Y.-C. et al. Foxp2 controls synaptic wiring of corticostriatal circuits and vocal communication by opposing Mef2c. Nat. Neurosci. 19, 1513–1522 (2016).

71. Dietrich, J.-B., Takemori, H., Grosch-Dirrig, S., Bertorello, A. & Zwiller, J. Cocaine induces the expression of MEF2C transcription factor in rat striatum through activation of SIK1 and phosphorylation of the histone deacetylase HDAC5. Synapse 66, 61–70 (2012).

72. Mega, T. et al. Zinc finger protein 521 antagonizes early B-cell factor 1 and modulates the B-lymphoid differentiation of primary hematopoietic progenitors. Cell Cycle 10, 2129–2139 (2011).

73. Zhang, Z. et al. Mutation of SLC35D3 Causes Metabolic Syndrome by Impairing Dopamine Signaling in Striatal D1 Neurons. PLOS Genet. 10, e1004124 (2014).

74. Russell, V. A. Reprint of ‘Neurobiology of animal models of attention-deficit hyperactivity disorder’. J. Neurosci. Methods 166, I–XIV (2007).

75. Ehrlich, M. E., Rosen, N. L., Kurihara, T., Shalaby, I. A. & Greengard, P. DARPP-32 development in the caudate nucleus is independent of afferent input from the substantia nigra. Brain Res. Dev. Brain Res. 54, 257–263 (1990).

76. Markowitz, J. E. et al. The Striatum Organizes 3D Behavior via Moment-to-Moment Action Selection. Cell 174, 44–58.e17 (2018).

77. Arama, J. et al. GABAA receptor activity shapes the formation of inhibitory synapses between developing medium spiny neurons. Front. Cell. Neurosci. 9, 290 (2015).

78. Arber, C. et al. Activin A directs striatal projection neuron differentiation of human pluripotent stem cells. Dev. Camb. Engl. 142, 1375–1386 (2015).

79. Faedo, A. et al. Differentiation of human telencephalic progenitor cells into MSNs by inducible expression of Gsx2 and Ebf1. Proc. Natl. Acad. Sci. U. S.A. 114, E1234–E1242 (2017).

80. Lin, L., Yuan, J., Sander, B. & Golas, M. M. In Vitro Differentiation of Human Neural Progenitor Cells Into Striatal GABAergic Neurons. Stem Cells Transl. Med. 4, 775–788 (2015).

81. Penrod, R. D., Campagna, J., Panneck, T., Preese, L. & Lanier, L. M. The presence of cortical neurons in striatal-cortical co-cultures alters the effects of dopamine and BDNF on medium spiny neuron dendritic development. Front. Cell. Neurosci. 9, 269 (2015).

82. Corbin, J. G. & Butt, S. J. B. Developmental mechanisms for the generation of telencephalic interneurons. Dev. Neurobiol. 71, 710–732 (2011).

83. Marin, O., Anderson, S. A. & Rubenstein, J. L. Origin and molecular specification of striatal interneurons. J. Neurosci. Off. J. Soc. Neurosci. 20, 6063–6076 (2000).

84. Dodson, P. D. et al. Distinct Developmental Origins Manifest in the Specialized Encoding of Movement by Adult Neurons of the External Globus Pallidus. Neuron 86, 501–513 (2015).

85. Nóbrega-Pereira, S. et al. Origin and molecular specification of globus pallidus neurons. J. Neurosci. Off. J. Soc. Neurosci. 30, 2824–2834 (2010).

86. Bielle, F. et al. Slit2 Activity in the Migration of Guidepost Neurons Shapes Thalamic Projections during Development and Evolution. Neuron 69, 1085–1098 (2011).

87. López-Bendito, G. et al. Tangential Neuronal Migration Controls Axon Guidance: A Role for Neuregulin-1 in Thalamocortical Axon Navigation. Cell 125, 127–142 (2006).

88. Tinterri, A. et al. Tangential migration of corridor guidepost neurons contributes to anxiety circuits. J. Comp. Neurol. 526, 397–411 (2018).

89. Waclaw, R. R., Ehrman, L. A., Pierani, A. & Campbell, K. Developmental Origin of the Neuronal Subtypes That Comprise the Amygdalar Fear Circuit in the Mouse. J. Neurosci. 30, 6944–6953 (2010).

90. Bupesh, M., Abellán, A. & Medina, L. Genetic and experimental evidence supports the continuum of the central extended amygdala and a mutiple embryonic origin of its principal neurons. J. Comp. Neurol. 519, 3507–3531 (2011).

91. Nadarajah, B. & Parnavelas, J. G. Modes of neuronal migration in the developing cerebral cortex. Nat. Rev. Neurosci. 3, 423–432 (2002).

92. Boller, S. & Grosschedl, R. The regulatory network of B-cell differentiation: a focused view of early B-cell factor 1 function. Immunol. Rev. 261, 102–115 (2014).

93. Demilly, A. et al. Coe genes are expressed in differentiating neurons in the central nervous system of protostomes. PloS One 6, e21213 (2011).

94. Dubois, L. & Vincent, A. The COE–Collier/Olf1/EBF–transcription factors: structural conservation and diversity of developmental functions. Mech. Dev. 108, 3–12 (2001).

95. Garcia-Dominguez, M. Ebf gene function is required for coupling neuronal differentiation and cell cycle exit. Development 130, 6013–6025 (2003).

96. Garel, S. et al. Family of Ebf/Olf-1-related genes potentially involved in neuronal differentiation and regional specification in the central nervous system. Dev. Dyn. Off. Publ. Am. Assoc. Anat. 210, 191–205 (1997).

97. Gyory, I. et al. Transcription factor Ebf1 regulates differentiation stage-specific signaling, proliferation, and survival of B cells. Genes Dev. 26, 668–682 (2012).

98. Kratsios, P., Stolfi, A., Levine, M. & Hobert, O. Coordinated regulation of cholinergic motor neuron traits through a conserved terminal selector gene. Nat. Neurosci. 15, 205–214 (2011).

99. de Taffin, M. et al. Genome-Wide Mapping of Collier In Vivo Binding Sites Highlights Its Hierarchical Position in Different Transcription Regulatory Networks. PloS One 10, e0133387 (2015).

100. Tursun, B., Patel, T., Kratsios, P. & Hobert, O. Direct conversion of C. elegans germ cells into specific neuron types. Science 331, 304–308 (2011).

101. Boller, S. et al. Pioneering Activity of the C-Terminal Domain of EBF1 Shapes the Chromatin Landscape for B Cell Programming. Immunity 44, 527–541 (2016).

102. Guilhamon, P. et al. Meta-analysis of IDH-mutant cancers identifies EBF1 as an interaction partner for TET2. Nat. Commun. 4, 2166 (2013).

103. Snyder-Keller, A. M. Development of striatal compartmentalization following pre-or postnatal dopamine depletion. J. Neurosci. 11, 810–821 (1991).

104. Lin, H. & Grosschedl, R. Failure of B-cell differentiation in mice lacking the transcription factor EBF. Nature 376, 263–267 (1995).

105. Lokmane, L. et al. Sensory Map Transfer to the Neocortex Relies on Pretarget Ordering of Thalamic Axons. Curr. Biol. 23, 810–816 (2013).

106. Xu, Q., Tam, M. & Anderson, S. A. Fate mapping Nkx2.1-lineage cells in the mouse telencephalon. J. Comp. Neurol. 506, 16–29 (2008).

107. Espinosa-Medina, I. et al. The sacral autonomic outflow is sympathetic. Science 354, 893–897 (2016).

108. Anders, S. & Huber, W. Differential expression analysis for sequence count data. Genome Biol. 11, R106 (2010).

